# *PRESENILIN 1* mutations causing early-onset familial Alzheimer’s disease or familial acne inversa differ in their effects on genes facilitating energy metabolism and signal transduction

**DOI:** 10.1101/2021.01.26.428321

**Authors:** Karissa Barthelson, Yang Dong, Morgan Newman, Michael Lardelli

## Abstract

**Background:** The most common cause of early-onset familial Alzheimer’s disease (EOfAD) is mutations in *PRESENILIN 1* (*PSEN1*) allowing production of mRNAs encoding full-length, but mutant, proteins. In contrast, a single known frameshift mutation in *PSEN1* causes familial acne inversa (fAI) without EOfAD. The molecular consequences of heterozygosity for these mutation types, and how they cause completely different diseases, remains largely unexplored.

**Objective:** To analyse brain transcriptomes of young adult zebrafish to identify similarities and differences in the effects of heterozygosity for *psen1* mutations causing EOfAD or fAI.

**Methods:** RNA sequencing was performed on mRNA isolated from the brains of a single family of 6-month-old zebrafish siblings either wild type or possessing a single, heterozygous EOfAD-like or fAI-like mutation in their endogenous *psen1* gene.

**Results:** Both mutations downregulate genes encoding ribosomal subunits, and upregulate genes involved in inflammation. Genes involved in energy metabolism appeared significantly affected only by the EOfAD-like mutation, while genes involved in Notch, Wnt and neurotrophin signalling pathways appeared significantly affected only by the fAI-like mutation. However, investigation of direct transcriptional targets of Notch signalling revealed possible increases in γ-secretase activity due to heterozygosity for either *psen1* mutation. Transcriptional adaptation due to the fAI-like frameshift mutation was evident.

**Conclusion:** We observed both similar and contrasting effects on brain transcriptomes of the heterozygous EOfAD-like and fAI-like mutations. The contrasting effects may illuminate how these mutation types cause distinct diseases.

## INTRODUCTION

Cases of Alzheimer’s disease (AD) can be classified by age of onset and mode of inheritance. Dominant mutations in a small number of genes cause AD with an age of onset younger than 65 years (early onset familial AD, EOfAD). On a population basis, around 60% of the mutations causing EOfAD occur in one gene, *PRESENILIN 1* (*PSEN1*) [1–4].

*PSEN1* encodes a multi-pass transmembrane protein resident in the endoplasmic reticulum, plasma membrane, endolysosomal pathway and other membranes [5, 6]. It has nine recognised transmembrane domains [7]. A tenth transmembrane domain may exist when PSEN1 protein is in its holoprotein state [7], before it undergoes autocatalytic endoproteolysis to form the active catalytic core of γ-secretase [8], an enzyme complex consisting of PSEN1 (or PSEN2), and the proteins NCSTN, PSENEN, and APH1A (or APH1B) [9, 10].

As a locus for genetic disease, *PSEN1* is truly remarkable both for the number of mutations found there, and the variety of diseases these mutations cause. Mutations have been found associated with Pick’s disease [11], dilated cardiomyopathy [12] and acne inversa [13]. However, over 300 mutations of *PSEN1* are known to cause EOfAD (www.alzforum.org/mutations/psen-1). In total, these mutations affect 161 codons of the gene. Remarkably, the mutations are widely distributed in the PSEN1 coding sequence, but are particularly common in the transmembrane domains. Only three regions of the PSEN1 protein are mostly devoid of EOfAD mutations: upstream of the first transmembrane domain; a large part of the “cytosolic loop domain” (cytosolic loop 3); and the last two thirds of the 9^th^ transmembrane domain together with the lumenal C-terminus (see **Figure 1**).

**Figure 1:**
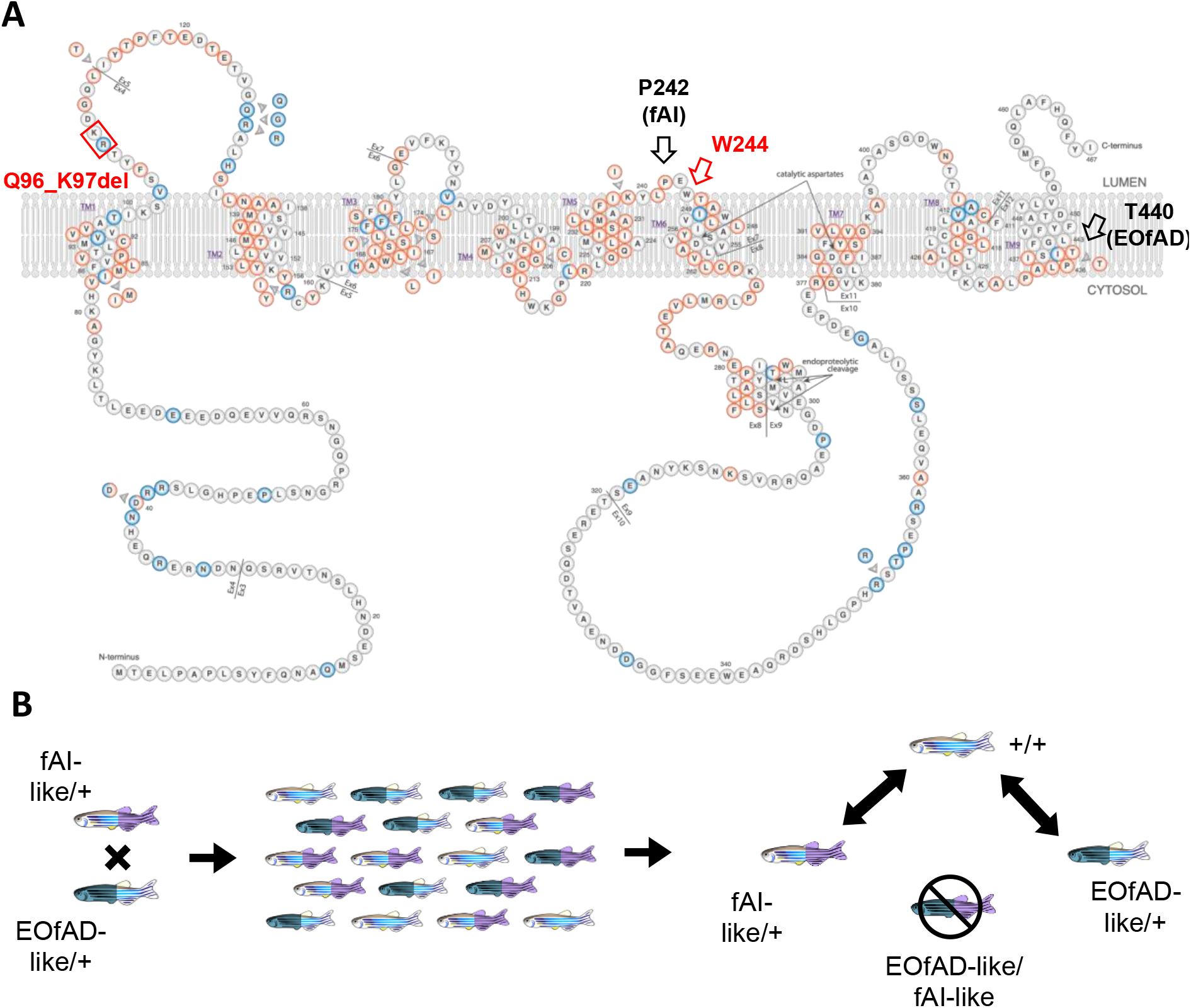
Experimental design. **A** Schematic of the human PSEN1 protein adapted from https://www.alzforum.org/mutations/psen-1 with permission from FBRI LLC (Copyright © 1996–2020 FBRI LLC. All Rights Reserved. Version 3.3 – 2020). Amino acid residues are colour-coded as to whether they are pathogenic for Alzheimer’s disease (red) or their pathogenicity is unclear (blue). The human mutation sites (P242 (fAI) and T440 (EOfAD) are indicated by black arrows. The site of the zebrafish W233fs-equivalent codon (W244) is shown by the red arrow. Note that the human T440 codon is equivalent to the zebrafish T428 codon. The residues equivalent to those deleted in the Q96_K97del mutation of zebrafish *psen1* analysed previously are indicated by a red box. **B** A fish heterozygous for the W233fs mutation (fAI-like/+) was mated with a fish heterozygous for the T428del mutation (EOfAD-like/+). The resulting family of fish contain genotypes fAI-like/+, EOfAD-like/+, EOfAD-like/fAI-like and their wild type siblings. The pairwise comparisons performed in the RNA-seq experiment are depicted. Since the EOfAD-like/fAI-like genotype is not representative of any human disease, it was not analysed. See online version for colour.

The most common outcome of mutation of a protein sequence is either no effect or a detrimental effect on the protein’s evolved activity. Only rarely are mutations selectively advantageous so that they enhance organismal survival and reproduction. The very large number of EOfAD-causative mutations in *PSEN1* and their wide distribution in the protein coding sequence is consistent with a loss-of-function. However, this cannot be a simple loss of γ-secretase activity, as EOfAD-causative mutations have never been found in the genes encoding the other components of γ-secretase complexes (other than less frequent mutations in the *PSEN1* homologous gene, *PSEN2*, reviewed in [14]). Also, an *in vitro* analysis of 138 EOfAD mutations of *PSEN1* published in 2017 by Sun et al. [15] found that approximately 10% of these mutations actually increased γ-secretase activity.

Currently, the most commonly discussed hypothetical mechanism addressing how EOfAD mutations of *PSEN1* cause disease is that these act through “qualitative changes” to γ-secretase cleavage of the AMYLOID β A4 PRECURSOR PROTEIN (AβPP) to alter the length distribution of the AMYLOID β (Aβ) peptides derived from it [16]. However, the comprehensive study of Sun et al. revealed no consistency in the effects of *PSEN1* EOfAD mutations on Aβ length distribution. The single consistent characteristic of all EOfAD mutations in both *PSEN1* and *PSEN2* is that these permit production of transcripts with coding sequences containing in-frame mutations, but terminated by the wild type stop codons (i.e. they still permit production of a full length protein). This phenomenon was first noted by De Strooper in 2007 [17] and described in detail by us in 2016 (the “reading frame preservation rule” [18]). The universality of this rule, and that it reflects a critical feature of the EOfAD-causative mechanism of *PSEN1* mutations, is shown by the fact that the P242LfsX11 frameshift mutation of *PSEN1* (hereafter referred to as P242fs) causes a completely different inherited disease, familial acne inversa (fAI, also known as hidradenitis suppurativa), *without* EOfAD [13]. (Recently, a frameshift mutation in *PSEN1*, H21PfsX2, was identified in an early-onset AD patient. However, whether the mutation is causative of EOfAD mutation is still uncertain [19]. Questionable additional claims of EOfAD-causative frameshift mutations in PSEN genes have been made and are reviewed in [18].) Critically, fAI can also be caused by mutations in *NCSTN* and *PSENEN* [13], strongly supporting that this disease is due to changes in γ-secretase activity.

Understanding the role of *PSENs* and their mutations is complicated by the partial functional redundancy shared by *PSEN1* and *PSEN2* and their complex molecular biology. For example, the PSEN1 holoprotein has been shown to have γ-secretase-independent activities required for normal lysosomal acidification [20], can form multimers [21–24], and may interact with the HIF1α protein [25–27] that is critical both for responses to hypoxia and for iron homeostasis (reviewed in [28]). Additionally, within γ-secretase complexes, the PSENs act to cleave at least 149 different substrates [29]. To simplify analysis, most previous investigation of PSEN activity has involved inactivation (knock-out) of *PSEN1* and/or *PSEN2* in cells or animals, and expression of only single forms of mutant PSEN (i.e. without simultaneous expression of wild type forms). Forced expression of *PSEN* genes is also usually at non-physiological levels which has unexpected regulatory feedback effects [30]. In humans, investigating *PSEN’s* role in AD at the molecular level is restricted to post-mortem brain tissues. However, these show substantially different patterns of gene expression compared to the brains of people with mild cognitive impairment (MCI) or age-matched healthy controls [31]. Since AD is thought to take decades to develop [32], we must understand the pathological effects of EOfAD mutations in young adult brains if we wish to identify preventative treatments. For this reason, we must model EOfAD mutations in animals.

The overwhelming majority of animal modelling of AD has utilised transgenic models favoured for their apparent, partial reproduction of Aβ histopathology and easily discernible cognitive disabilities. However, the relationship between Aβ histopathology and cognitive change in these models is questionable [33]. Additionally, the most detailed form of molecular phenotyping currently available, transcriptome analysis, shows little consistency between the disturbed brain gene transcription of various transgenic models and limited concordance between them and human sporadic AD brain transcriptomes [34]. “Knock-in” mouse models of single EOfAD mutations (e.g. [35, 36]) make the fewest assumptions regarding the pathological mechanism(s) of AD and most closely replicate the human EOfAD genetic state (i.e. incorporating a single, dominant, endogenous mutation in the heterozygous state). However, the brain transcriptomes of these mice have never been analysed, and interest in them waned due to their lack of Aβ histopathology and mild cognitive effects.

Analysis of mouse brain transcriptomes is complicated by strong effects on gene expression of sex [37, 38] and, potentially, litter-of-origin (i.e. due to environmental and genotypic variation) (K. Barthelson, unpublished results). In contrast, zebrafish brain transcriptomes show only subtle influences of sex, and very large numbers of siblings can be generated from single mating event, alleviating potential litter-of-origin issues [39–43]. In 2014, our laboratory began a program of creating knock-in models of EOfAD-like (and non-EOfAD-like) mutations in the zebrafish genes orthologous to *PSEN1, PSEN2*, and *SORL1*. In 2019 we began publishing the results of transcriptome analyses of the young adult brains of these fish [39–46] as we attempt to establish what effect(s) all the EOfAD mutations have in common (and differentiate them from the non-EOfAD-like mutations).

Our previous analyses of an EOfAD-like mutation in the zebrafish *psen1* gene, Q96_K97del, revealed very significant effects on the expression of genes involved in mitochondrial function, lysosomal acidification, and iron homeostasis [45, 46]. Although Q96_K97del follows the reading-frame preservation rule of EOfAD mutations in *PSEN1*, it is not an exact equivalent of any human EOfAD-causative mutation. Consequently, in this study we aimed to generate an additional, exactly equivalent, model of a human *PSEN1* EOfAD mutation. For technical reasons, the T428 codon of zebrafish *psen1* (equivalent to the T440 codon of human PSEN1) was predicted to be readily targetable using CRISPR-Cas9 technology, and we subsequently deleted this codon in the zebrafish gene. This generated a zebrafish model of the human EOfAD mutation *PSEN1^T440del^* [47]. This mutation was identified in a Japanese man classified as displaying a mixed dementia phenotype (variant AD with spastic paraparesis, Parkinson’s disease and dementia with Lewy bodies). Also, to understand how reading-frame preserving and frameshift mutations can cause completely different diseases we generated a frameshift mutation in zebrafish *psen1, psen1^W233fs^*, very similar to the fAI-causative P242fs mutation of human *PSEN1*. We then performed an RNA-seq analysis with high read depth and large sample numbers to compare the brain transcriptomes of fish from a single family of young adult siblings heterozygous for either mutation or wild type. We observed subtle, and mostly distinct, effects of the two mutations. In particular, changes in the fAI-like brain transcriptomes implied significant effects on Notch, Wnt, neurotrophin, and Toll-like receptor signalling, while changes in the EOfAD-like brain transcriptomes implied effects on oxidative phosphorylation similar to those previously seen for EOfAD-like mutations in *psen1* [45], *psen2* [40, 43], and *sorl1* [39, 41].

## MATERIALS AND METHODS

### Zebrafish husbandry and animal ethics

All zebrafish (Tübingen strain) used in this study were maintained in a recirculating water system on a 14 hour light/10 hour dark cycle, fed dry food in the morning and live brine shrimp in the afternoon. All zebrafish work was conducted under the auspices of the Animal Ethics Committee (permit numbers S-2017-089 and S-2017-073) and the Institutional Biosafety Committee of the University of Adelaide.

### CRISPR-Cas9 genome editing

To mutate zebrafish *psen1*, we used the Alt-R^®^ CRISPR-Cas9 system (Integrated DNA Technologies, Coralville, IA, USA). To generate the T428del mutation (EOfAD-like) in exon 11 of *psen1*, we used a custom-designed crRNA recognising the sequence 5’ CTCCCCATCTCCATAACCTT 3’ and a PAM of CGG. For the W233fs mutation (fAI-like), the crRNA was designed to recognise the sequence 5’ GATGAGCCATGCGGTCCACT 3’ in exon 6 of *psen1*, with a PAM sequence of CGG. We aimed to generate exact equivalents of the human P242fs mutation causing fAI, and the T440del mutation causing EOfAD by homology directed repair (HDR). For the P242fs mutation, we used a plasmid DNA template as described in [48] (synthesised by Biomatik, Kitchener, Ontario, Canada). For the T440del mutation, we used an antisense, asymmetric single-stranded oligonucleotide with phosphorothioate modifications (synthesised by Merck, Kenilworth, NJ, USA) as described in [49] (HDR template DNA sequences are given in **Supplementary File 1.**)

Each crRNA was annealed with an equal amount of Alt-R^®^ CRISPR-Cas9 tracrRNA (IDT) in nuclease free duplex buffer (IDT) by heating at 95°C for 5 minutes, then allowed to cool to room temperature, giving sgRNA solutions of 33 μM (assuming complete heteroduplex formation of the RNA molecules). Then, 1 μL of the sgRNA solution was incubated with 1 μL of Alt-R^®^ S.p.Cas9 Nuclease 3NLS (IDT) at 64 μM at 37°C for 10 minutes to form ribonucleoprotein (RNP) complexes. The final concentration for the linear ssODN for the T428del mutation was 1 μM, and the final concentration of the plasmid DNA for the W233fs mutation was 25 ng/μL. Approximately 2-5 nL of RNP complexes in solution with the respective template DNAs were injected into Tübingen strain zebrafish embryos at the one cell stage. The procedures followed for testing of the mutagenesis efficiencies of CRISPR-Cas9 systems using allele-specific polymerase chain reactions and T7 endonuclease I assays, and the breeding strategy to isolate the mutations of interest, are described in [41, 50].

### RNA-seq raw data generation and processing

We performed RNA-seq on a family of zebrafish as described in **Figure 1**. Total RNA (with genomic DNA depleted by DNaseI treatment) was isolated from the brains of n = 4 fish per genotype and sex as described in [41]. Then, 500 ng of total RNA (RINe > 9) was delivered to the South Australian Genomics Centre (SAGC, Adelaide, Australia) for polyA+ library construction (with unique molecular identifiers (UMIs)) and RNA-sequencing using the Illumina Novaseq S1 2×100 SE platform.

The raw fastq files from SAGC were provided as 100 bp paired end reads as well as an index file containing the UMIs for each read (over two Novaseq lanes which were subsequently merged). The merged raw data was processed using a developed Nextflow [51] RNA-seq workflow (see https://github.com/sagc-bioinformatics/sagc-rnaseq-nf). Briefly, UMIs were added to headers of each read using *fastp* (v0.20.1). Alignment of the reads to the zebrafish genome (GRCz11, Ensembl release 98) was performed using *STAR* (v2.5.3a). Then, reads which contained the same UMI (i.e. PCR duplicates) were deduplicated using the *dedup* function of *umi_tools* (version 1.0.1). Finally, the gene-level counts matrix was generated using *featureCounts* from the *Subread* (version 2.0.1) package.

### Differential gene expression

Statistical analysis of the RNA-seq data was performed using *R* [52]. Since lowly expressed genes are considered uninformative for differential expression analysis, we omitted genes with less than 0.1 counts per million (CPM) (following the 10/minimum library size in millions rule described in [53]). Library sizes after omitting the lowly expressed genes ranged between 61 and 110 million reads. These were normalised using the trimmed mean of M-values (TMM) method [54]. To test for differential expression of genes due to heterozygosity for the T428del or W233fs mutation, we used a generalised linear model and likelihood ratio tests using *edgeR* [55, 56]. A design matrix was specified with the wild type genotype as the intercept, and the T428del/+ and W233fs/+ genotypes as the coefficients. We considered a gene to be differentially expressed (DE) due to each *psen1* mutant genotype if the FDR adjusted p-value was less than 0.05.

### Enrichment analysis

We tested for over-representation of gene ontology (GO) terms within the DE gene lists using *goseq* [57], using the average transcript length per gene to calculate the probability weighting function (PWF). We considered a GO term to be significantly over-represented within the DE gene lists relative to all detectable genes in the RNA-seq experiment if the FDR-adjusted p-value generated by *goseq* was less than 0.05.

We also performed enrichment analysis on the entire list of detectable genes by calculating the harmonic mean p-value from the raw p-values calculated from *fry* [58], *camera* [59] and *fgsea* [60, 61] as described in [41], To test for changes to gene expression in a broad range of biological processes, we used the KEGG [62] gene sets obtained from MSigDB [63] using the *msigdbr* package [64]. We also used *msigdbr* to obtain gene sets which contain genes that show changed expression in response to changes in the Notch signalling pathway (*NGUYEN_NOTCH1_TARGETS_UP* and *NGUYEN_NOTCH1_TARGETS_DN, NOTCH_DN.V1_UP, NOTCH_DN.V1_DN* and *RYAN_MANTLE_CELL_LYMPHOMA_NOTCH_DIRECT_UP*). We also tested for evidence of possible iron dyshomeostasis using gene sets containing genes encoding transcripts which contain iron-responsive elements in their untranslated regions (described in [45]).

### Comparison of the T428del and Q96_K97del mutations in psen1

Isolation of the zebrafish Q96_K97del mutation in zebrafish *psen1* and analysis of its effects on zebrafish brain transcriptomes have been described previously [45, 46]. That dataset is comprised of brain RNA-seq data for fish heterozygous for the Q96_K97del mutation and their wild type siblings, at 6 months old (young adult) and 24 months old (aged), and under normoxia or hypoxia treatment (n = 4 fish per genotype, age and treatment). In the analysis presented here, we performed enrichment analysis using the methods described above on the entire dataset, but presented the results for the pairwise comparison between 6-month-old Q96_K97del/+ fish and wild type fish under normoxia.

To obtain a broader comparison on the effects of the Q96_K97del and T428del mutations, we performed adaptive, elastic-net sparse PCA (AES-PCA) [65] as implemented in the *pathwayPCA* package [66]. For this analysis, we utilised the HALLMARK [67] gene sets from MSigDB to generate the pathway collection. The pathway principal components (PCs) were calculated only on the gene expression data from samples heterozygous for an EOfAD-like mutation (Q96_K97del or T428del) and their wild type siblings under normoxia at 6 months of age. Then, the categorical effect of genotype was tested for association with the pathway PCs using a permutation-based regression model as described in [66].

### Data availability

The paired end fastq files and the output of *featureCounts* have been deposited in GEO under accession number GSE164466. Code to reproduce this analysis can be found at https://github.com/karissa-b/psen1_EOfAD_fAI_6m_RNA-seq.

## RESULTS

### Generation of an EOfAD-like and a fAI-like mutation in zebrafish psen1

An unsolved puzzle regarding the dominant EOfAD mutations of human *PSEN1* (and *PSEN2*) is why these are consistently found to permit production of transcripts in which the reading frame is preserved, while heterozygosity for mutations causing frameshifts (or deleting the genes) does not cause EOfAD. To investigate this quandary in an *in vivo* model, we initially aimed to generate mutations in zebrafish *psen1* which would be exact equivalents of the T440del and P242fs mutations using homology directed repair (HDR). While screening for the desired mutations, we identified the mutations W233fs and T428del, both likely generated by the non-homologous end joining (NHEJ) pathway of DNA repair. T428del is a 3 nucleotide deletion which, nevertheless, produces a protein-level change exactly equivalent to that observed for the human T440del mutation. Hereafter, for simplicity, we refer to the T428del mutation as “EOfAD-like”. W233fs is an indel mutation causing a frameshift in the second codon downstream of the zebrafish *psen1* proline codon equivalent to human *PSEN1* codon P242. This change is still within the short third lumenal loop of the Psen1 protein (see **Figure 1**). Assuming no effect on splicing, the frameshift mutation results in a premature stop codon 36 codons downstream of W233 (**Figure S1** in **Supplementary File 2**). Hereafter, we refer to this mutation as “fAI-like”. The alignment of the wild type and mutant sequences in humans and zebrafish is shown in **Figure 2**.

**Figure 2:**
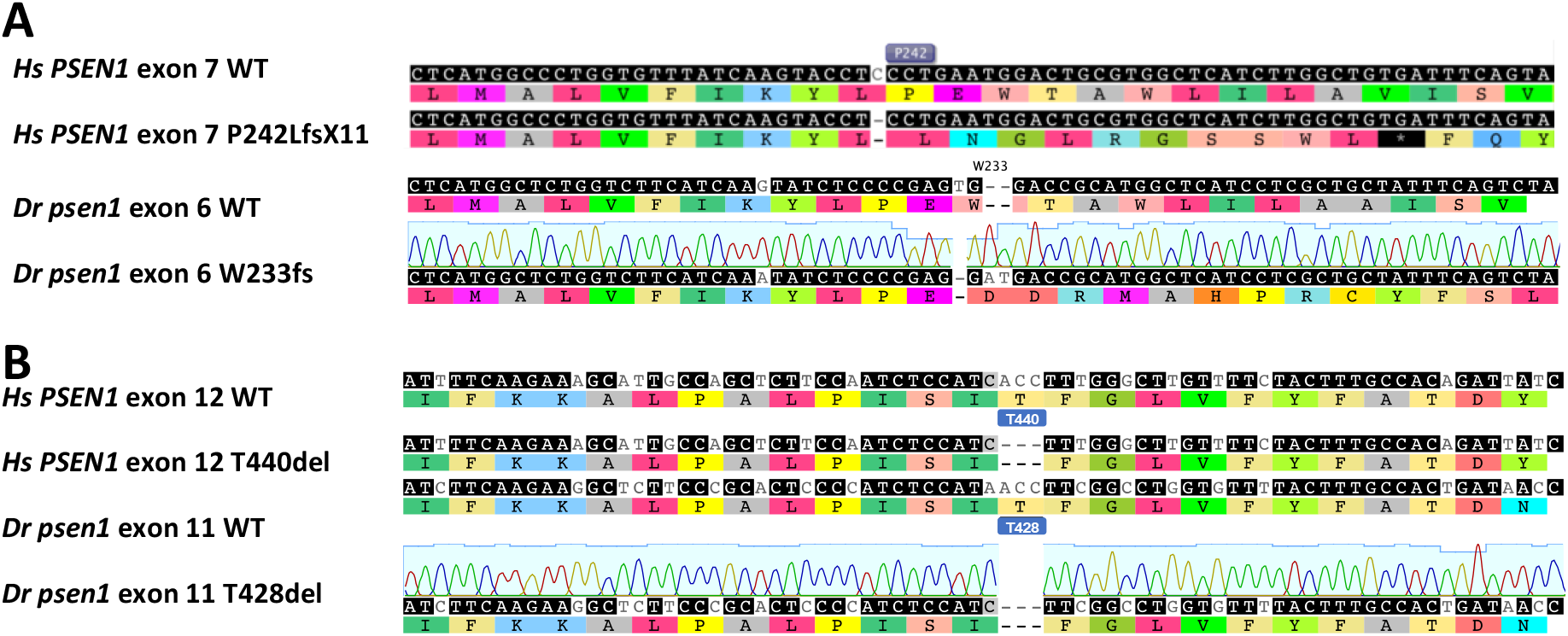
**A** Alignment of a region of wild type human (*Hs*) *PSEN1* exon 7, and the same region containing the human P242fs (P242LfsX11) mutation, and the equivalent zebrafish (*Dr*) *psen1* exon 6 wild type and W233fs sequences. **B** Alignment of a region of wild type human (*Hs*) *PSEN1* exon 12, and the same region containing the human T440del mutation, and the equivalent zebrafish (*Dr*) *psen1* exon 11 wild type and T428del sequences.

Heterozygosity or homozygosity for neither the EOfAD-like nor the fAI-like mutation produces any obvious morphological defects. However, this is unsurprising considering that rare examples of humans homozygous for EOfAD mutations are known [68, 69] and loss of *PSEN1* γ-secretase activity is, apparently, compatible with viability in zebrafish [70] and rats [71] (although not in mice [72]).

### Transcriptome analysis

To investigate global changes to the brain transcriptome due to heterozygosity for the EOfAD-like or fAI-like mutations in *psen1*, we performed mRNA-seq on a family of fish as described in **Figure 1**. The family of sibling fish were raised together in a single tank, thereby reducing sources genetic and environmental variation between individuals and allowing subtle changes to the transcriptome to be detected with minimal confounding effects.

To begin our exploration of the similarities and differences between the brain transcriptomes of the mutant and wild type fish, we first performed principal component analysis (PCA) on the gene level, log transformed counts per million (logCPM) of the zebrafish RNA-seq samples. A plot of principal component 1 (PC1) against PC2 did not show distinct clustering of samples by genotype or sex, indicating that these variables do not result in stark changes to the brain transcriptome. This is consistent with our previous observations of EOfAD-like mutations in other genes [39–41]. However, some separation of the EOfAD-like and fAI-like samples is observed across PC2, indicating distinct, but subtle, differences between these transcriptomes. Notably, the majority of the variation in this dataset is not captured until PC6 (**Figure 3**).

**Figure 3:**
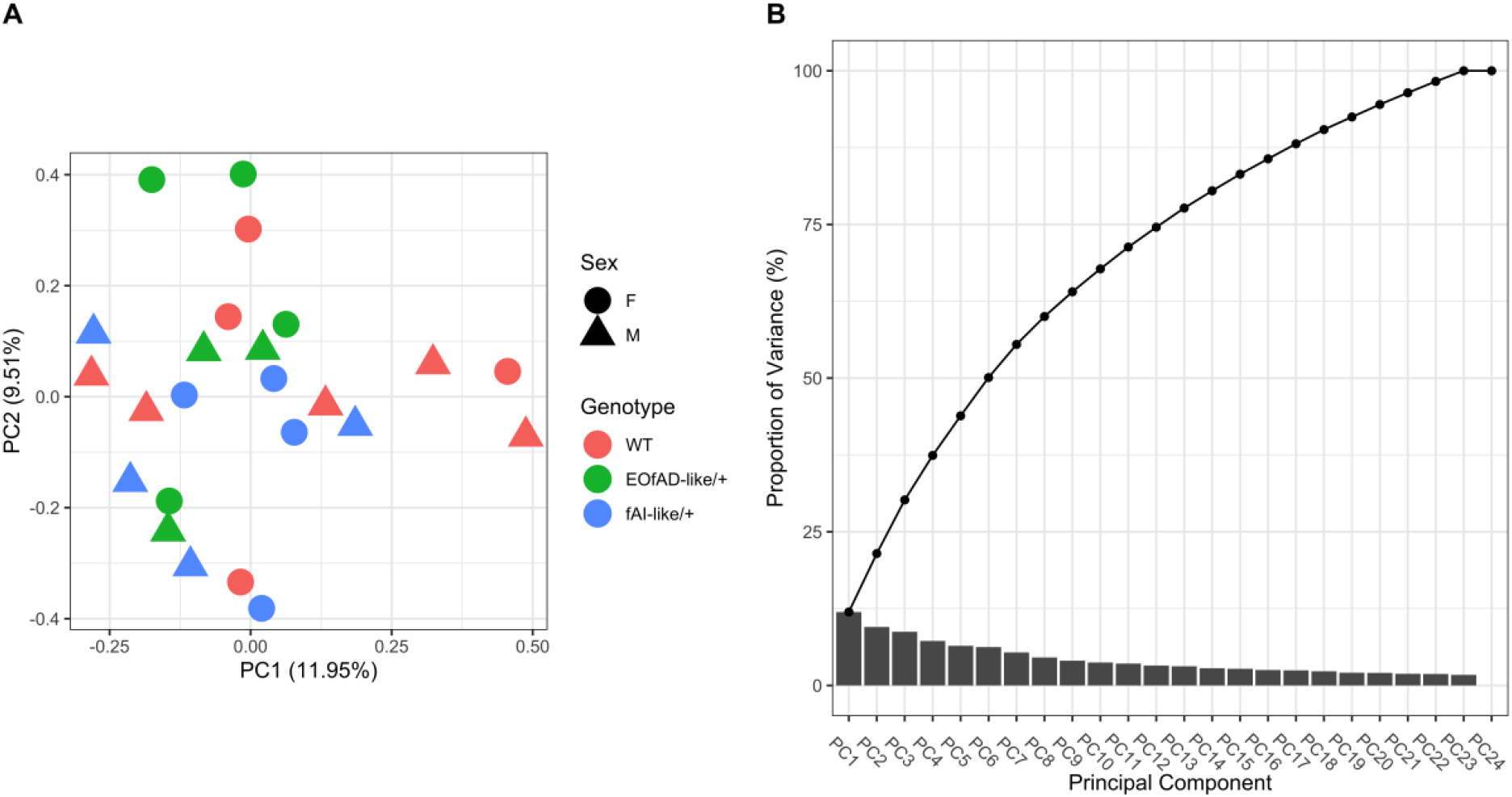
Principal component (PC) analysis of the gene expression values for the RNA-seq experiment. **A** PC1 plotted against PC2 for each sample. Each point represents a sample and is coloured according to *psen1* genotype. Female (F) samples appear as circles and male (M) samples appear as triangles. **B** Scree plot indicating the variance explained by each principal component. The points joined by lines indicate the cumulative variance explained by each PC. See online version for colour.

### Heterozygosity for the EOfAD-like or fAI-like mutations of psen1 causes only subtle effects on gene expression

Which genes are dysregulated due to heterozygosity for the EOfAD-like or the fAI-like mutations? To address this question, we performed differential gene expression analysis using a generalised linear model and likelihood ratio tests with *edgeR*. We observed statistical evidence for 13 genes as significantly differentially expressed (DE) due to heterozygosity for the EOfAD-like mutation, and 5 genes due to the fAI-like mutation (**Figure 4, Supplementary Table 1**). Notably, *psen1* was the most significantly DE gene due to heterozygosity for the fAI-like mutation (logFC = −0.8, FDR = 1.33e-78), consistent with the observation that frame-shift mutations commonly induce nonsense-mediated mRNA decay when they result in premature stop codons (reviewed in [73]). Total levels of *psen1* transcripts were unchanged in EOfAD-like/+ brains (logFC = −0.0065, FDR = 1, **Figure S7** in **Supplementary File 2**). No DE genes were found to be shared between the comparisons of either form of heterozygous mutant to wild type, or were found to be significantly overrepresented by any gene ontology (GO) terms by *goseq* (for the top 5 most significantly over-represented GO terms in each comparison, see **Tables S1** and **S2** in **Supplementary File 2**). This is not unexpected due to the relatively low number of significantly DE genes detected.

**Figure 4:**
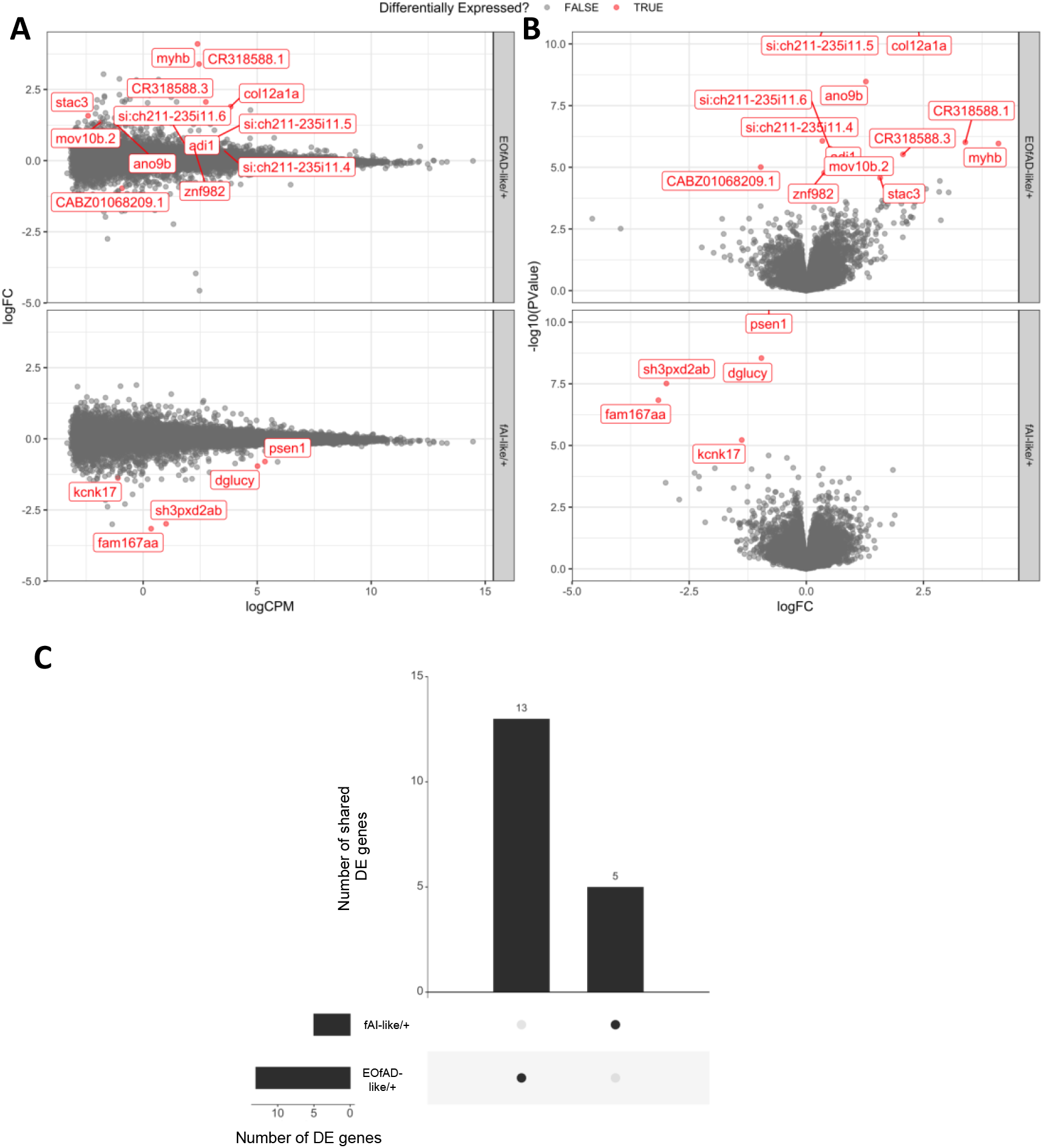
Differential expression analysis. **A** Mean-difference (MD) plots and **B** volcano plots of changes to gene expression in EOfAD-like/+ and fAI-like/+ mutant zebrafish brains. Note that the limits of the y-axis in **B** are restrained to between 0 and 10 for visualisation purposes. **C** Upset plot indicating the low number of genes which are significantly differentially expressed (DE) in either comparison. See online version for colour.

### Significant differences in gene expression between the EOfAD-like and fAI-like mutants can be detected at the pathway level

Since very few DE genes were detected in each comparison of heterozygous *psen1* mutant fish to their wild type siblings, we performed enrichment analysis on all detectable genes in the RNA-seq experiment. Our method, inspired by the *EGSEA* framework [74], involves calculation of the harmonic mean p-value [75] from the raw p-values of three different rank-based gene set testing methods: *fry* [58], *camera* [59] and *GSEA* [60, 61]. Unlike *EGSEA*, we use the harmonic mean p-value to combine the raw p-values, as the harmonic mean p-value has been specifically shown to be robust for combining dependent p-values [75]. We performed enrichment testing using the KEGG gene sets (describing 186 biological pathways and processes) to obtain information on changes to activities for these pathways. We also tested for evidence of iron dyshomeostasis using our recently defined sets of genes containing iron-responsive elements (IREs) in the untranslated regions of their mRNAs [45]. We observed statistical evidence for 7 KEGG gene sets as significantly altered by heterozygosity for the EOfAD-like mutation and 11 KEGG gene sets as significantly altered by heterozygosity for the fAI-like mutation (**Figure 5,** full results of the raw p-values from each algorithm as well as the harmonic mean p-value can be found in **Supplementary Table 2**).

**Figure 5:**
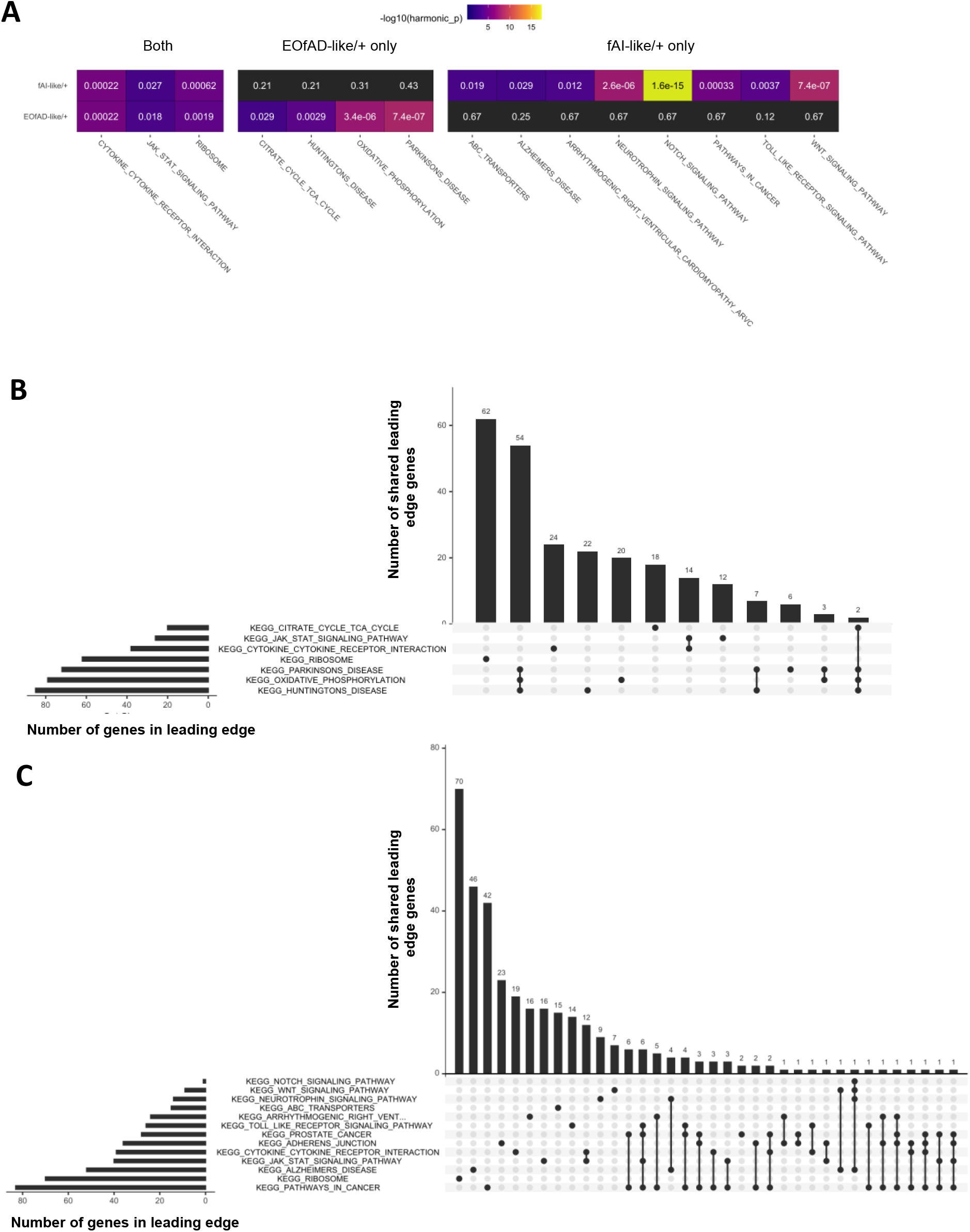
**A** KEGG gene sets with FDR-adjusted harmonic mean p-values of < 0.05 in *psen1* EOfAD-like/+ and fAI-like/+ mutant brains. The colour of the cells indicates the level of significance (brighter colour indicates greater statistical significance, while dark grey indicates the FDR-adjusted harmonic mean p-value > 0.05). The number within each cell is the FDR-adjusted harmonic mean p-value. **B** Upset plot indicating the overlap of leading edge genes from the fgsea algorithm which drive the enrichment of gene sets significantly altered in EOfAD-like/+ and **C** fAI-like/+ brains. See online version for colour.

Gene sets significantly altered in the brains of both forms of heterozygous mutant included the KEGG gene sets for cytokine receptor interactions, Jak/Stat signalling, and encoding the components of the ribosomal subunits. Inspection of the leading edge genes (which can be interpreted as the core genes driving the enrichment of a gene set) showed that similar genes were driving the enrichment of the gene sets for cytokine receptor interactions and Jak/Stat signalling. Similar genes were also driving the enrichment of the *KEGG_RIBOSOME* gene set in both heterozygous mutants. However, the magnitude of the logFC was greater in the fAI-like/+ samples, suggesting a stronger effect (**Figure S4-S6** in **Supplementary File 2**). Gene sets which were only altered significantly by heterozygosity for the EOfAD-like mutation were involved in energy metabolism (*KEGG_PARKINSONS_DISEASE, KEGG_OXIDATIVE_PHOSPHORYLATION, and KEGG_CITRATE_CYCLE_TCA_CYCLE*). Notably, the KEGG gene sets for Parkinson’s disease, Huntington’s disease, and for oxidative phosphorylation, share 55 leading-edge genes, implying that their enrichment is driven by, essentially, the same gene expression signal (**Figure 5**). Conversely, the 10 KEGG gene sets found to be altered significantly by heterozygosity for the fAI-like mutation appear to be driven mostly by distinct gene expression signals. No IRE gene sets were observed statistically to be altered in the brains of either mutant, suggesting that iron homeostasis is unaffected (at least at 6 months of age). The changes to expression of genes within the KEGG gene sets are likely not due to broad changes in cell-type proportions in the zebrafish brain samples, since the expression of marker genes of neurons, astrocytes, oligodendrocytes and microglia was similar in all samples (**Figure S13** in **Supplementary File 2**).

### The EOfAD-like and fAI-like mutations alter expression of Notch signalling genes

Notch signalling plays a critical role in many cell differentiation events and is dependent on PSEN’s γ-secretase activity. Disturbance of Notch signalling due to decreased γ-secretase activity has been suggested to contribute to the changes in skin histology of fAI, as Notch signalling is required for normal epidermal maintenance ([76–78] and reviewed by [79]). However, fAI has not been reported as associated with EOfAD, despite that the T440del mutant form of *PSEN1* appears to have little intrinsic γ-secretase activity [15]. The expression of genes involved in the KEGG gene set for the Notch signalling pathway was observed to be highly significantly altered in the brains of fAI-like/+ mutants, but not of EOfAD/+ mutants, implying that y-secretase activity might only be affected significantly by the frameshift, fAI-like mutation (**Figure 5**). However, inspection of the logFC of genes in the *KEGG_NOTCH_SIGNALING_PATHWAY* gene set revealed similar patterns of changes to gene expression in both mutants (**Figure 6**). Upregulation of the genes encoding the Notch and Delta receptors is observed in both mutants compared to wild type. In fAI-like/+ brains, we observe downregulation of the downstream transcriptional targets of the Notch intracellular domain (NICD), implying decreased Notch signalling (and, likely, reduced y-secretase activity). Genes encoding repressors of Notch signalling are observed to be upregulated (i.e. *dyl* and *numb*), reinforcing this interpretation.

**Figure 6:**
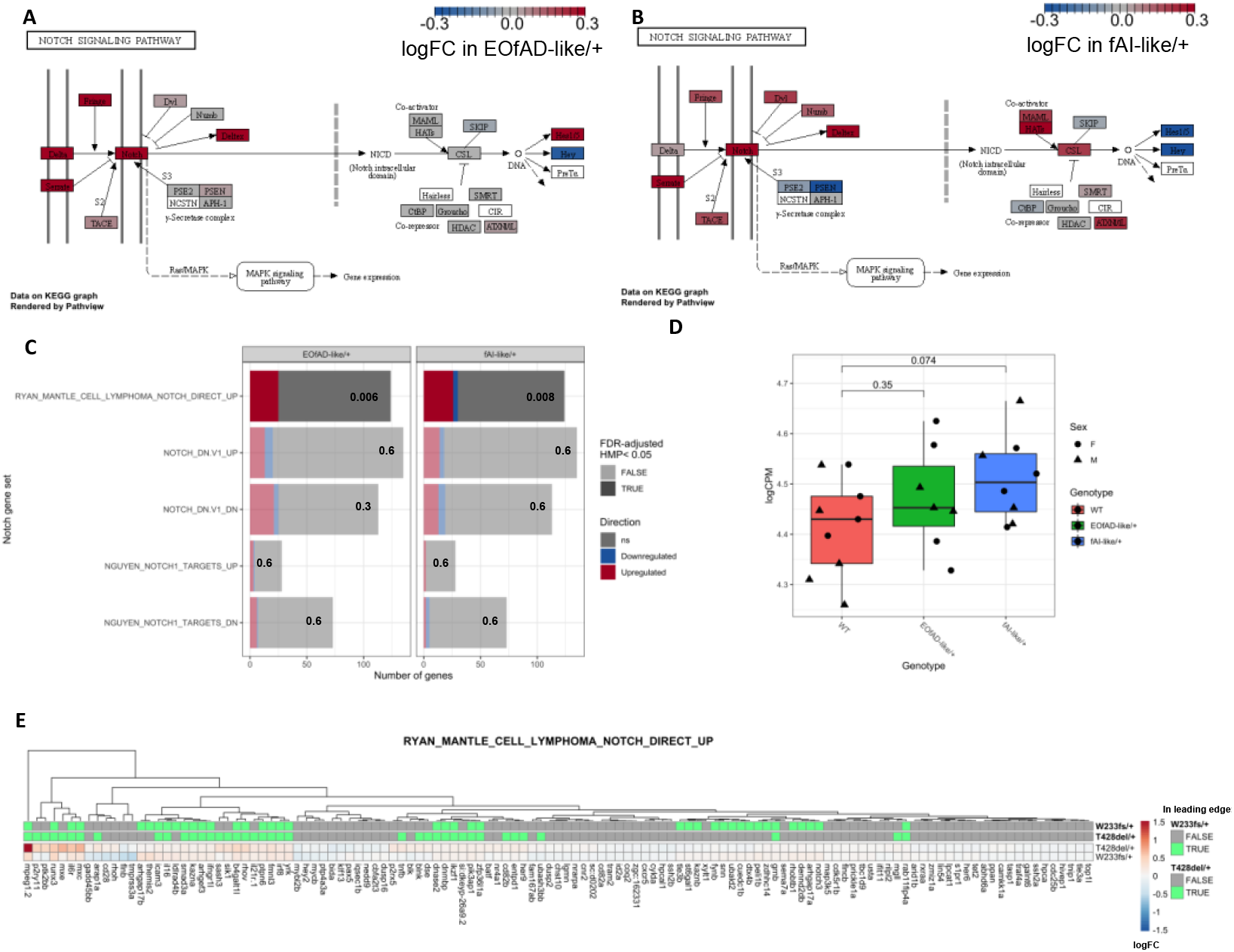
**A** Pathview [85] visualisation of the changes to gene expression in the *KEGG_NOTCH_SIGNALING_PATHWAY* gene set in EOfAD-like/+ mutants and **B** fAI-like/+ mutants. **C** The proportion of genes with increased expression (red, z > √2) and decreased expression (blue, z < √2) in MSigDB gene sets for Notch signalling in EOfAD-like/+ and fAI-like/+ mutant brains. Gene sets which contained a FDR-adjusted harmonic mean p-value (HMP) < 0.05 appear less transparent. The FDR adjust p-values are also listed on the bars. **D** The expression of *psen2* is trending towards upregulation, particularly in fAI-like/+ mutants. Here, p-values were determined by Student’s unpaired t-tests. **E** Heatmap indicating the logFC values for genes in the *RYAN_MANTLE_CELL_LYMPHOMA_NOTCH_DIRECT_UP* gene set. Genes are clustered based on their Euclidean distance, and are labelled with green if they appear in the leading edge of the *fgsea* algorithm for each comparison of a *psen1* heterozygous mutant with wild type. See online version for colour.

Since the KEGG gene set for Notch signalling only contains two genes that are direct transcriptional targets of the NICD, we investigated further whether Notch signalling is perturbed in both mutants by analysis of gene sets from MSigDB containing information on genes responsive to Notch signalling in different cell lines: *NGUYEN_NOTCH1_TARGETS_UP; NGUYEN_NOTCH1_TARGETS_DN; NOTCH_DN.V1_UP; NOTCH_DN.V1_DN;* and *RYAN_MANTLE_CELL_LYMPHOMA_NOTCH_DIRECT_UP*. The *NGUYEN_NOTCH1_TARGETS_UP* and *_DOWN* gene sets consist of genes which have been observed as up- or downregulated respectively in response to a constitutively active Notch receptor in keratinocytes [80]. *The NOTCH_DN.V1_UP* and *_DN* gene sets contain genes which are up- and down-regulated respectively in response to treatment with the *γ*-secretase inhibitor DAPT in a T-cell acute lymphoblastic leukemia (T-ALL) cell line [81]. The *RYAN_MANTLE_CELL_LYMPHOMA_NOTCH_DIRECT_UP* gene set contains genes showing both increased expression upon rapid activation of Notch signalling by washout of the *γ-* secretase inhibitor compound E, and evidence for a NICD binding site in the promotor by ChIP-seq, in mantle cell lymphoma cell lines [82]. (Note that there is no equivalent “*RYAN_MANTLE_CELL_LYMPHOMA*” gene set representing genes downregulated in response to Notch signalling.) Of these 5 gene sets, statistical support was found only for changes to the expression of genes in the *RYAN_MANTLE_CELL_LYMPHOMA_NOTCH_DIRECT_UP* gene set, and this was found for both the EOfAD-like (p=0.006) and the fAI-like (p=0.008) mutants. The leading edge genes were mostly observed to be upregulated, which supports increases in Notch signalling (implying increased y-secretase activity). Transcriptional adaptation (previously known as “genetic compensation”) might contribute to the apparent increase in Notch signalling in the frameshift, fAI-like/+ mutant brains via upregulated expression of the *psen1*-homologous gene, *psen2* [83, 84]. Although no statistically significant differences in expression were observed for *psen2* in the differential expression test using *edgeR* (see **Supplementary Table 1**), a trend towards upregulation in the fAI-like/+ mutants was observed following a simple Student’s t-test (p=0.074, **Figure 6D**). El-Brolosy et al. [83] showed that the wild type allele of a mutated gene can also be upregulated by transcriptional adaptation (where the mutation causes nonsense-medicated decay, NMD, of mutant transcripts). Inspection of the number of reads aligning to the W233 mutation site across samples indicates that the expression of the wild type *psen1* allele in fAI-like/+ brains appears to be greater than 50% of the expression of the wild type *psen1* allele in wild type brains (p = 0.006), providing further evidence for transcriptional adaptation due to the fAI-like mutation (**Figure S8** in **Supplementary File 2**).

Together, these results suggest that Notch signalling and, by implication, γ-secretase activity, may be enhanced in *psen1* mutant brains. However, future biochemical assays should be performed to confirm this prediction.

### The EOfAD-like mutation T428del has a milder phenotype than the previously studied Q96_K97del EOfAD-like mutation of psen1

The T428del mutation of *psen1* is the first identified zebrafish mutation exactly equivalent, (at the protein sequence level), to a characterised human EOfAD mutation. Therefore, we sought to assess the consistency of its effects with those of a previously studied EOfAD-like mutation, Q96_K97del, and to identify cellular processes affected in common by the two mutations. The Q96_K97del mutation deletes two codons in the sequence encoding the first lumenal loop of the Psen1 protein (see **Figure 1**). Comparison of transcriptomes from the 6-month-old brains of Q96_K97del/+ and wild type siblings previously predicted changes to expression of genes involved in energy metabolism, iron homeostasis and lysosomal acidification [45, 46]. To compare which cellular processes are affected by heterozygosity for the Q96_K97del mutation or the T428del mutation, we first performed enrichment analysis on the RNA-seq data previously generated by our analysis of zebrafish heterozygous for the Q96_K97del mutation relative to their wild type siblings. Here, we observed that heterozygosity for the Q96_K97del mutation results in significant alterations in 7 KEGG gene sets (at 6 months of age during normoxia, Figure 7A). We also found statistical evidence for altered expression of genes possessing IREs in their 3’ UTRs (see *IRE3_ALL* in **Figure 7A**), consistent with our previous finding using a different method of enrichment analysis [45]. Gene sets affected in common between the two EOfAD-like mutations in *psen1* are involved in energy metabolism and protein translation (**Figure 7A**). The expression of genes involved in protein degradation, and of genes containing IREs in the 3’ UTRs of their transcripts, appeared significantly altered only by the Q96_K97del mutation (**Figure 7A**).

**Figure 7:**
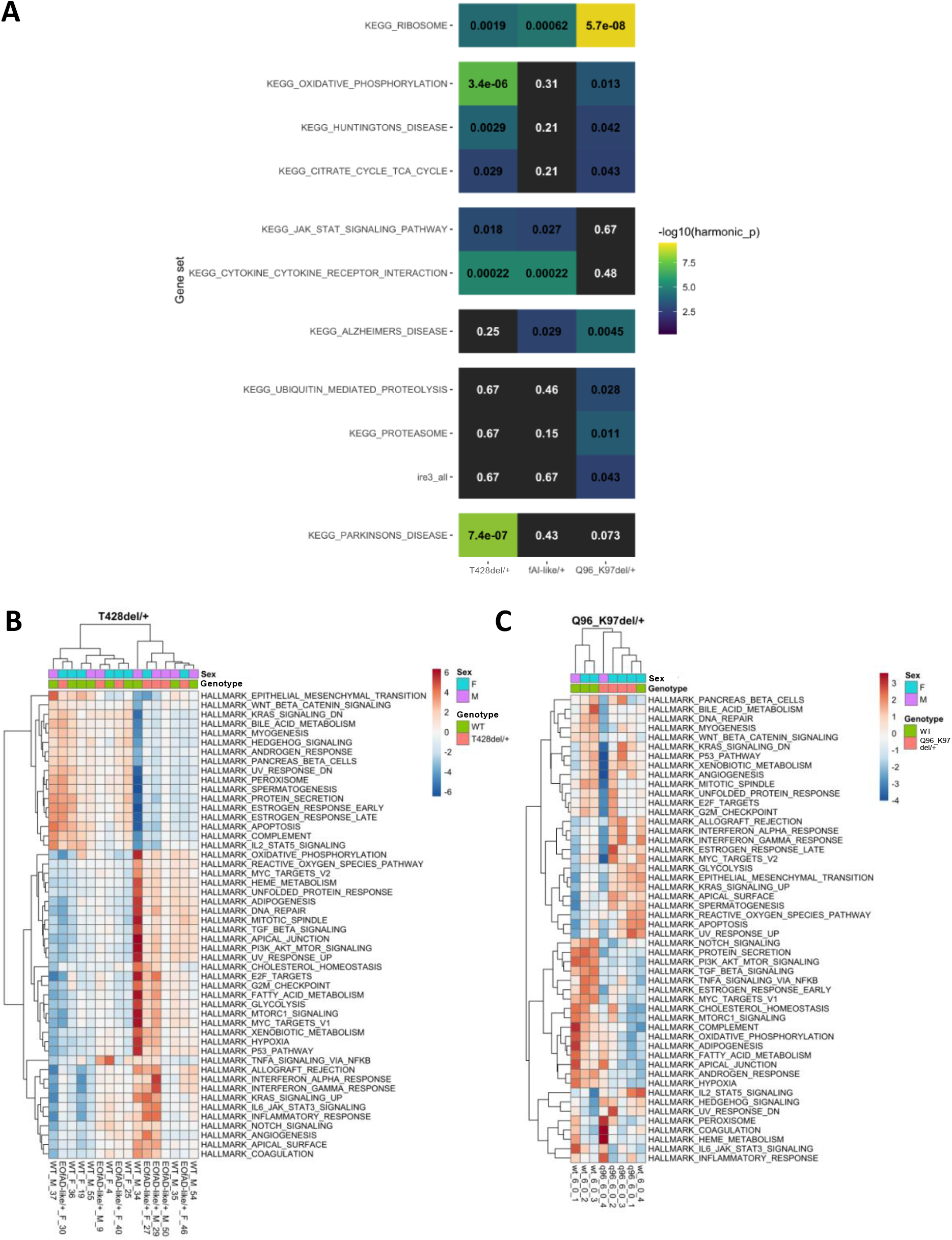
**A** Comparison of KEGG and IRE gene sets significantly altered by the EOfAD-like mutations T428del and Q96_K97del in 6-month-old zebrafish brains. Each cell is coloured according to statistical significance, and the FDR-adjusted harmonic mean p-value is shown. Gene sets not significantly altered (FDR adjusted harmonic mean p-value > 0.05) in a comparison between a *psen1* mutant zebrafish with their respective wild type siblings appear grey. **B** Principal component 1 (PC1) values for the HALLMARK gene sets as calculated by AES-PCA, clustered based on their Euclidean distance in T428del/+ samples relative to their wild type siblings. **C** PC1 values for the HALLMARK gene sets as calculated by AES-PCA clustered based on their Euclidean distance in Q96_K97del/+ samples relative to their wild type siblings at 6 months of age under normal oxygen conditions. See online version for colour.

We also compared the effects of the two EOfAD-like mutations using adaptive, elastic-net sparse PCA (AES-PCA) as implemented in the *pathwayPCA* package [66]. AES-PCA allows reduction of data dimensionality and for the overall activity of predefined gene sets to be observed in a sample-specific manner [65]. To obtain a global view of the changes to gene expression between the two *psen1* EOfAD-like mutations over the two datasets, we utilised the HALLMARK gene sets that encompass 50 distinct biological processes (rather than the 186 KEGG gene sets that share many genes).

The latent variables estimated by AES-PCA for the HALLMARK gene sets (i.e. the first principal components) in each dataset did not show any significant association with *psen1* genotype, suggesting that changes to gene expression (measured over entire brains) are too subtle to be detected as statistically significant using this method. However, clustering of the calculated PC1 values by AES-PCA for each HALLMARK gene set in each sample and dataset revealed that samples in the Q96_K97del dataset clustered mostly according to genotype (one wild type sample did not follow the trend), supporting that heterozygosity for the Q96_K97del mutation does result in marked effects on gene expression for the HALLMARK gene sets. Conversely, clustering of PC1 values in the T428del dataset resulted in two distinct clusters of samples. However, samples did not group by genotype over the two clusters to the same extent as seen for the Q96_K97del dataset. Intriguingly, the Q96_K97del dataset had less sample numbers per genotype (n = 4), and did not have as great sequencing depth as the current RNA-seq experiment. Therefore, this supports that heterozygosity for the Q96_K97del mutation has more consistent (more severe) effects on young adult brain transcriptomes than heterozygosity for the T428del mutation (**Figure 7 B,C**).

## DISCUSSION

In this study, we exploited transcriptome analysis of whole brains of young adult zebrafish siblings, to detect differences in molecular state between the brains of fish heterozygous for an EOfAD-like mutation or an fAI-like mutation of *psen1* compared to wild type *in vivo*. The subtlety of the effects observed is consistent with that EOfAD is, despite its designation as “early-onset”, a disease affecting people overwhelmingly at ages older than 30 years [86]. The person reported to carry the T440del mutation of *PSEN1* showed cognitive decline at 41 years [47]. (Overall, EOfAD mutations in *PSEN1* show a median survival to onset of 45 years [86]). In contrast, at 6 months of age, zebrafish are only recently sexually mature. Nevertheless, since AD is thought to take decades to develop [32], it is these subtle, early changes that we must target therapeutically if we wish to arrest the pathological processes driving the progression to AD. As seen in all our previous analyses of EOfAD-like mutations [39–41, 43, 45], changes in expression of genes involved in oxidative phosphorylation were identified as significant. However, this was not the case for the fAI-like, frameshift mutation. Therefore, oxidative phosphorylation changes appear to be an early signature cellular stress of EOfAD. Changes to mitochondrial function have been observed in heterozygous *PSEN1* mutant astrocytes [87], homozygous *PSEN1* mutant neurons [88], and in neurons differentiated from human induced pluripotent stem cells (hIPSCs) from sporadic AD patients [89], supporting our findings. However, such changes are not always observed [90], possibly due to issues of experimental reproducibility between laboratories when working with hIPSCs [91].

The EOfAD-like mutation also caused very statistically significant changes in the *KEGG_PARKINSONS_DISEASE* gene set (that shares many genes with *KEGG_OXIDATIVE_PHOSPHORYLATION*) and the person carrying the *PSEN1^T440del^* mutation modelled by zebrafish *psen1^T428del^* initially showed symptoms of early onset parkinsonism at 34 years of age before those of cognitive decline at 41 years [47].

In contrast to this EOfAD-like mutation, the fAI-like mutation apparently caused very statistically significant changes in Notch signalling and changes in other signal transduction pathways such as those involving Wnt and neurotrophins, as might be expected from changes in γ-secretase activity. Also notable was enrichment for the *KEGG_TOLL_LIKE_RECEPTOR_SIGNALLING_PATHWAY* gene set since acne inversa is a chronic inflammatory skin disorder and, in humans, increased expression of Toll-like receptor 2 has been noted in acne inversa lesions [92].

Both the EOfAD-like and fAI-like mutations caused very statistically significant changes in the gene sets *KEGG_CYTOKINE_CYTOKINE_RECEPTOR_INTERACTION* and *KEGG_RIBOSOME*. The former gene set reflects that both mutations appear to affect inflammation that is a characteristic of the pathologies of both EOfAD [93] and fAI (reviewed in [94]). Like oxidative phosphorylation, we have also observed effects on ribosomal protein genes sets for every EOfAD-like mutation we have studied [39–41, 43]. This may be due protein synthesis consuming a large proportion of cells’ energy budgets [95] and requiring amino acid precursors that can be sourced from lysosomes. Recently, Bordi et al [96] noted that mTOR is highly activated in fibroblasts from people with Down syndrome (DS, trisomy 21). DS individuals commonly develop EOfAD due to overexpression of the AβPP gene (that is resident on human chromosome 21). The consequent increased expression of AβPP’s β-CTF/C99 fragment (generated by β-secretase cleavage of AβPP without γ-secretase cleavage) affects endolysosomal pathway acidification [97] in a similar manner to EOfAD mutations of *PSEN1* [98]. The mTOR protein is localised at lysosomes in the mTORC1 and mTORC2 protein complexes (reviewed in [99, 100]) and monitors the energy and nutrient status of cells (reviewed in [101]). It is important for regulating ribosomal activity, partly by regulating transcription of ribosome components (reviewed in [102, 103]). Therefore, one explanation for the consistent enrichment for transcripts of the *KEGG_RIBOSOME* gene set we see in EOfAD mutant brains may be mTOR activation due to effects on lysosomal acidification and/or the energy status of cells. We did not observe any significant changes to the expression of genes involved in the mTOR signalling pathway in the analyses described in this paper (the FDR-adjusted harmonic mean p-value for the *KEGG_MTOR_SIGNALING_PATHWAY* was 0.7 for each comparison of the *psen1* mutant fish to their wild type siblings, see **Supplementary Table 2**). However, these changes could be undetectably subtle in young adult brains and/or occurring at the protein level and therefore not observable in bulk RNA-seq data. (Statistically significant enrichment for genes in the *HALLMARK_PI3K_AKT_MTOR_SIGNALING* gene set was seen previously for the normoxic, 6-month-old brains of fish heterozygous for the more severe EOfAD-like mutation Q96_K97del when compared to wild type siblings [45].)

While the brain transcriptome alterations caused by the EOfAD-like and fAI-like mutations are subtle (as illustrated by the lack of tight clustering of samples in the principal component analysis in **Figure 3,** and the low number of significantly differentially expressed genes), we are reassured in their overall veracity by their similarity to the results of a parallel analysis of sibling brain transcriptomes from 6-month-old zebrafish heterozygous for either a frameshift or a frame-preserving mutation in the zebrafish *psen2* gene relative to wild type [40]. In that similarly structured (but less statistically powered) experiment, only the frameshift mutation significantly affected the *KEGG_NOTCH_SIGNALLING* gene set while only the frame-preserving, EOfAD-like mutation significantly affected the *KEGG_OXIDATIVE_PHOSPHORYLATION* gene set. Both *psen2* genotypes affected the *KEGG_RIBOSOME* gene set, but in overall opposite directions (the frameshift mutation largely upregulated these genes while the frame-preserving mutation did the opposite).

Transcriptome analysis can reveal a great deal of data on differences in gene transcript levels between different genotypes or treatments. However, interpreting changes in cellular state from this information is not straight forward. Are any changes seen direct molecular effects of a mutation/treatment (e.g. the direct, downstream effects of a change in γ-secretase activity) or homeostatic responses as cells/tissues adjust their internal states to promote survival? For example, in the *KEGG_NOTCH_SIGNALING_PATHWAY* gene set shown in **Figure 6B**, more pathway components are upregulated than are downregulated. However, the direct transcriptional targets of Notch signalling (*her4.2* and *heyl*) are downregulated, as might be expected from reduced expression of wild type, catalytically-competent Psen1 protein. The upregulation of other components of the pathway may represent homeostatic responses attempting to restore normal levels of Notch signalling.

Only two Notch downstream transcriptional target genes are described in the *KEGG_NOTCH_SIGNALING_PATHWAY* gene set. Therefore, in an effort to assess more generally the effects of the EOfAD-like and fAI-like mutations on γ-secretase activity, we also analysed additional sets of genes previously identified (in various systems) as direct transcriptional targets of γ-secretase-dependent signalling. One of these sets, encompassing genes identified as Notch signalling targets by both γ-secretase inhibitor responses and binding of the Notch intracellular domain to chromatin, revealed apparent upregulation of Notch signalling in both the EOfAD-like and fAI-like heterozygous mutant brains relative to wild type siblings. In both EOfAD-like/+ and fAI-like/+ mutant brains, the most highly ranked genes in terms of differential expression tended to be upregulated (although some highly ranked genes were downregulated in fAI-like/+ brains). The idea that putative low levels of a form of Psen1 protein truncated in the third lumenal loop domain could increase Notch signalling is not unexpected, as we previously observed an implied upregulation of Notch signalling in zebrafish embryos with forced expression of the fAI-causative P242Lfs allele of human *PSEN1* [104]. However, a widespread assumption within AD research is that EOfAD-like mutations of *PSEN1* decrease γ-secretase cleavage of AβPP [105], possibly through a dominant negative mechanism [21]. This assumption conflicts with the observation of Sun et al. [15] that approximately 10% of the 138 EOfAD mutations of human *PSEN1* they studied actually increased γ-secretase cleavage of AβPP’s β-CTF/C99 fragment (in experiments examining the activities of the mutant proteins in isolation from wild type protein). Zhou et al. [106] also observed increased γ-secretase activity (cleavage of AβPP’s β-CTF/C99 fragment) due to an EOfAD mutation of *PSEN1* (S365A, this replicated Sun et al.’s finding for this mutation).

It is important to note that mutations of *PSEN1* need not cause similar effects on Notch and AβPP cleavage [104, 107, 108]. The transmembrane domains of the Notch receptor and the APP’s C99 fragment have different conformations [109]. Therefore, changes in the conformation of PSEN1 within γ-secretase due to a mutation may differentially affect Notch and C99 cleavage. Indeed, in our previously mentioned study of forced expression of the human *PSEN1^P242Lfs^* allele in zebrafish embryos, increased apparent Notch signalling was observed without change in AβPP processing [104]. Conversely, Zhang et al. [108] showed that transgenic expression of an EOfAD mutation S169del in *PSEN1* under the control of the *Thy1* brain specific promotor altered the processing of AβPP *in vivo* without affecting Notch signalling. Notably, both of these studies did not use the *PSEN1* gene’s own promoter to express mutant forms of this gene, and so the effects seen may be distorted by gene/protein over-expression.

Unfortunately, the direct transcriptional targets of the intracellular domain of AβPP (AICD) have not been characterised to the same extent as those of NICD (reviewed in [110]). This constrains transcriptome analysis for detection of differential effects on γ-secretase cleavage of AβPP caused by the EOfAD-like and fAI-like mutations. Future work should include further investigation of how these mutations effect γ-secretase cleavage of AβPP *in vivo*.

In conclusion, we have performed the first direct comparison of an EOfAD-like and a fAI-like mutation of *presenilin 1* in an *in vivo* model. Both forms of mutation cause apparent changes in inflammation, downregulate expression of genes encoding the components of the ribosome subunits, and potentially affect γ-secretase activity as supported by altered expression of Notch signalling pathway transcriptional target genes. We see that changes to mitochondrial function are a specific, common characteristic of EOfAD-like mutations while the fAI-like mutation specifically affects important signal transduction pathways. These differential effects on brain transcriptomes give insight into how reading-frame preserving mutations in *PSEN1* cause EOfAD while frameshift mutations do not.

## CONFLICT OF INTEREST/DISCLOSURE STATEMENT

The authors have no financial or non-financial competing interests to declare.

## ACKNOWLEDGEMENTS

The authors would like to thank Dr. Nhi Hin for providing the Q96_K97del gene expression values and the zebrafish IRE gene sets. We also would like to thank Dr. Jimmy Breen for his assistance with using the Nextflow pipeline. The authors thank Dr Giuseppe Verdile for critical reading of the manuscript.

This work was supported with supercomputing resources provided by the Phoenix HPC service at the University of Adelaide and by grants GNT1061006 and GNT1126422 from the National Health and Medical Research Council of Australia (NHMRC). KB was supported by an Australian Government Research Training Program Scholarship and by funds from the Carthew Family Charity Trust. YD was supported by an Adelaide Graduate Research Scholarship from the University of Adelaide. MN was supported by funds from the grants listed above. ML is an academic employee of the University of Adelaide.

## Notes

### Competing Interest Statement

The authors have declared no competing interest.

### Summary of Updates

Corrected a mistake in Figure 2

