## Supplemental File 1 for "*PRESENILIN 1* mutations causing early-onset familial Alzheimer’s disease or familial acne inversa differ in their effects on genes facilitating energy metabolism and signal transduction"

### Template DNA sequences for attempted HDR

**Generation of the P242fs mutation in zebrafish *psen1.***

The following sequence was supplied in the pBluescript II SK(+)-Amp vector. The sequences for the sgRNA are indicated in red, PAM sites are highlighted in blue, and the site of the mutation site is highlighted in yellow.

Template plasmid insert sequence 5’ – 3’:

GATGAGCCATGCGGTCCACTCGGTATACATTATTAAAAAAAGCACACAATACTGTCTGTTTCTTTTGCATAGACCTGAATAATGAATACACTTTTACAGTGCTGTAAAATACAAGCCTACACACAACCTTTTTTTTTTTGCTATTTTGCTGAAAACTAAAAATTTTTAGGGCAAGTTTGACATTTGCATGGAATTGCTCAGTATTAACGTGATTTCCTGCCTCTTAACGATGTTTCTGTTCCCTCCAGGGAAGTGTTCAAGACGTATAACGTGGCGATGGATTACTTCACGCTGGCGTTGATCATCTGGAACTTCGGTGTGGTGGGAATGATCTGCATCCACTGGAAGGGGCCGCTGCGGCTCCAGCAGGCCTATCTGATCATGATCAGCGCTCTCATGGCTCTGGTCTTCATCAAGTATCTCCTGAATGGACTGCGTGGCTCATCCTGGCTGTGATTTCAGTCTACGGTCAGTCAGTCTGCAAACAGAGCAACACATTCACTTCTGTGTACTGATAGGTTTATTGTTCTGTAATGCTGCTTTGGAACAATAATTAGCCTGAAATGAATTGAATAATTTCAAAAACACAGCCCCTTAAATGTCTTTCCTGCTCCTGTGTAGATCTTCTGGCAGTGTTGTGTCCGAAAGGCCCTCTGCGAATCCTTGTGGAAACAGCTCAAGAGAGGAATGAGGCCATTTTCCCAGCGCTCATCTACTCCTGTAAGAAAAACCCCTCAGCACTTCCATCTTCTCAGACTGTGAATGTCCATCTGTTTAATTGTGTTATCGTCACGTTTCTATGAAATTTCTCCTTCGATTTGTCAGCTACGATGGTGTGGCTCTTCAATATGGCGGACCGAGTGGACCGCATGGCTCATC

**Generation of the T428del mutation in zebrafish *psen1.***

A linear, single-stranded oligonucleotide (ssODN) with phosphorothioated (*) ends. The PAM is indicated in blue.

ssODN sequence 5’ – 3’:

C*G*ACCTCTCTATATGTAGAACTGATGGACGGCCAGCTGGTCCATGAACGGCCGCACGAGGTTATCAGTGGCAAAGTAAAACACCAGGCCGAATATGGAGATGGGGAGTGCGGGAAGAG*C*C
