## Supplemental File 2 for "*PRESENILIN 1* mutations causing early-onset familial Alzheimer’s disease or familial acne inversa differ in their effects on genes facilitating energy metabolism and signal transduction"

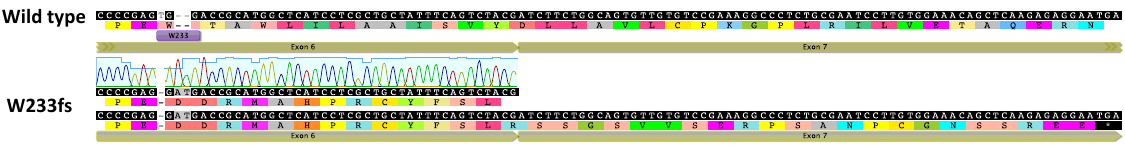

**Figure S1:** Alignment of the a region of *psen1* cDNA (ENSDART00000149864.3) including a subset of exons 6 and 7. The top sequence is the WT sequence, which is followed by a subset of the Sanger sequencing (from genomic DNA) of exon 6 from W233fs homozygous mutants. The bottom sequence is the predicted coding sequences of exon 6 and exon 7 assuming splicing is unaffected by the W233fs mutation. 36 novel amino acids are generated from this mutation followed by a premature termination codon.

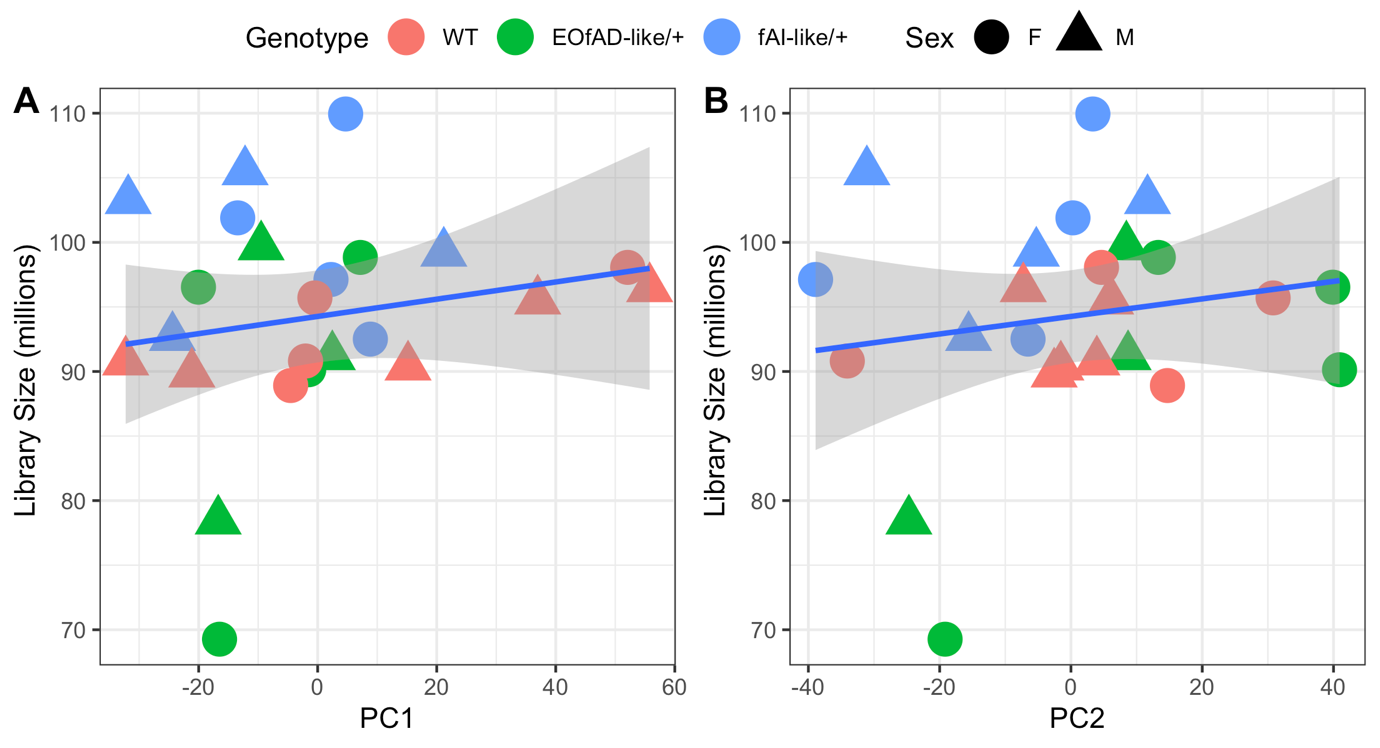

**Figure S2:** RNA-seq library size does correlate with **A** principal component 1 (PC1) or **B** PC2 after weighted trimmed mean of M-values (TMM) normalisation. Blue linear regression lines are shown with standard error shading.

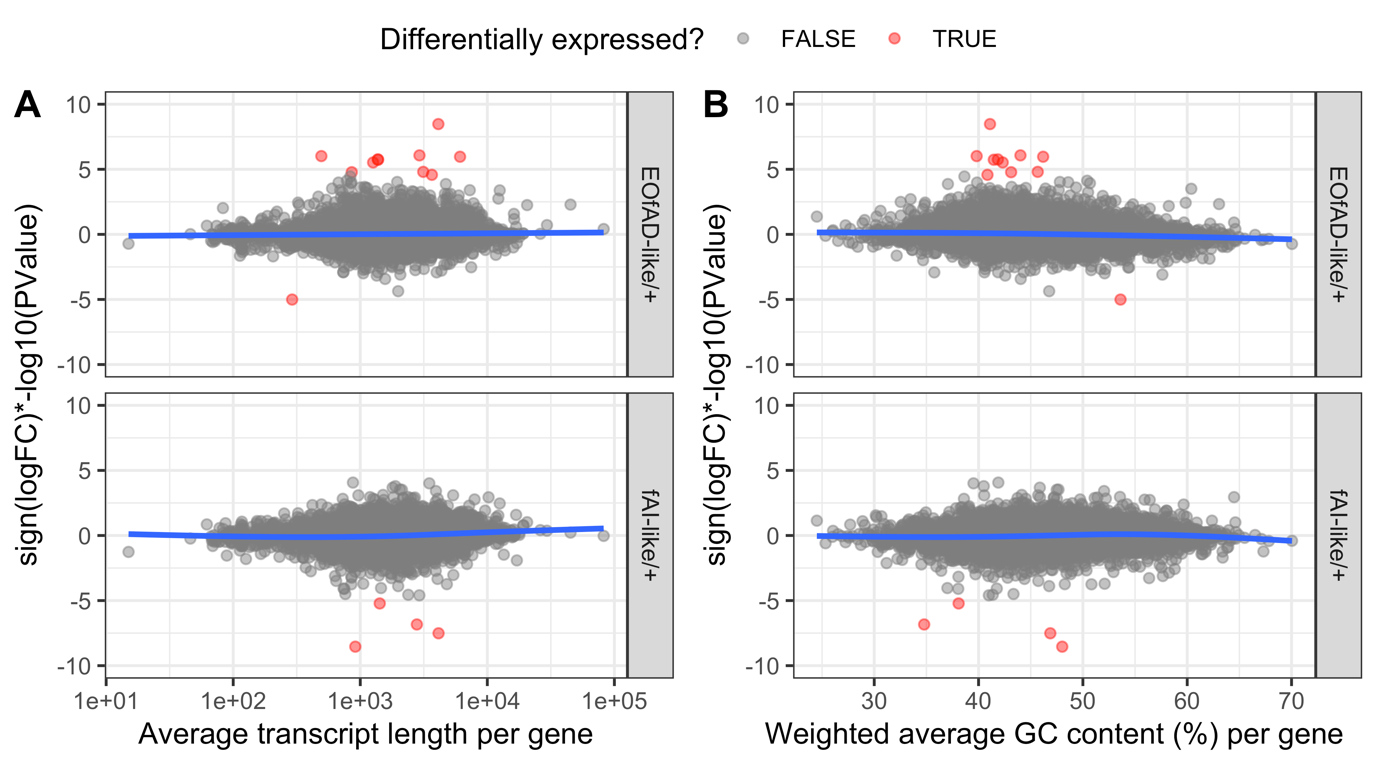

**Figure S3:** No observed bias for differential expression of genes due to **A** average transcript length or **B** weighted (per transcript length) GC content per gene. A ranking statistic was calculated for each gene. Blue lines indicating generalised additive model fits lie mostly around 0. The y axis in each graph is constrained to between -10 and 10 for visualisation purposes.

| **Table S1:** Top 5 GO terms over-represented in genes significantly differentially expressed (DE) in EOfAD-like/+ relative to wild type brains. | | | | |
| --- | --- | --- | --- | --- |
| **GO term** | **p.val** | **Number of DE genes in category** | **Number of genes in category** | **FDR** |
| GO SKELETAL MUSCLE CONTRACTION | 3.5e-05 | 2 | 32 | 0.35 |
| GO MULTICELLULAR ORGANISMAL MOVEMENT | 6.8e-05 | 2 | 44 | 0.35 |
| GO STRIATED MUSCLE CONTRACTION | 7.8e-04 | 2 | 147 | 1 |
| GO CALCIUM ACTIVATED  PHOSPHOLIPID SCRAMBLING | 1.2e-03 | 1 | 4 | 1 |
| GO L METHIONINE SALVAGE FROM  METHYLTHIOADENOSINE | 1.6e-03 | 1 | 5 | 1 |
| **Table S2:** Top 5 GO terms over-represented in genes significantly differentially expressed (DE) in fAI-like/+ relative to wild type brains | | | | |
| **GO term** | **p.val** | **Number of DE genes in category** | **Number of genes in category** | **FDR** |
| GO GAMMA SECRETASE COMPLEX | 0.0009 | 1 | 4 | 1 |
| GO ASPARTIC ENDOPEPTIDASE ACTIVITY INTRAMEMBRANE CLEAVING | 0.001 | 1 | 5 | 1 |
| GO NOTCH RECEPTOR PROCESSING LIGAND DEPENDENT | 0.001 | 1 | 5 | 1 |
| GO CATION CHANNEL ACTIVITY | 0.001 | 2 | 276 | 1 |
| GO REGULATION OF L GLUTAMATE IMPORT ACROSS PLASMA MEMBRANE | 0.001 | 1 | 6 | 1 |

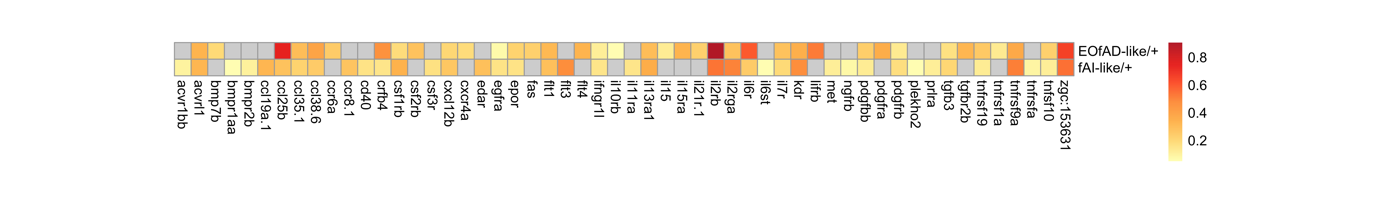

**Figure S4:** Similar genes are driving the enrichment of the *KEGG_CYTOKINE_CYTOKINE_RECEPTOR_INTERACTION* in the same direction. Colour of the heatmap indicates the logFC of the leading edge genes from the fgsea algorithm. Grey indicates that the gene was not found in the leading edge.

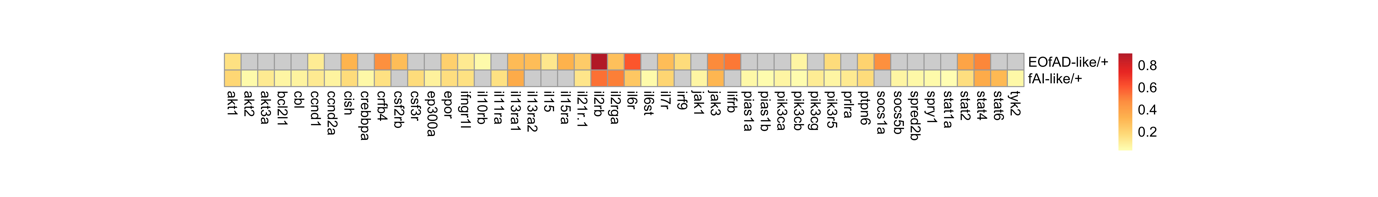

**Figure S5:** Mostly similar genes are driving the enrichment of the *KEGG_JAK_STAT_SIGNALING_PATHWAY* gene set in the same direction. Colour of the heatmap indicates the logFC of the leading edge genes. Grey indicates that the gene was not found in the leading edge in the fgsea algorithm.

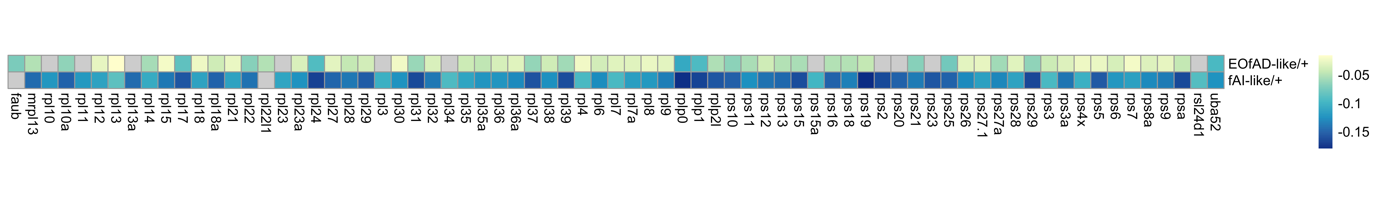

**Figure S6:** Similar genes are driving the enrichment of the *KEGG_RIBOSOME* gene set in the same direction. Colour of the heatmap indicates the logFC of the leading edge genes. Grey indicates that the gene was not found in the leading edge in the fgsea algorithm.

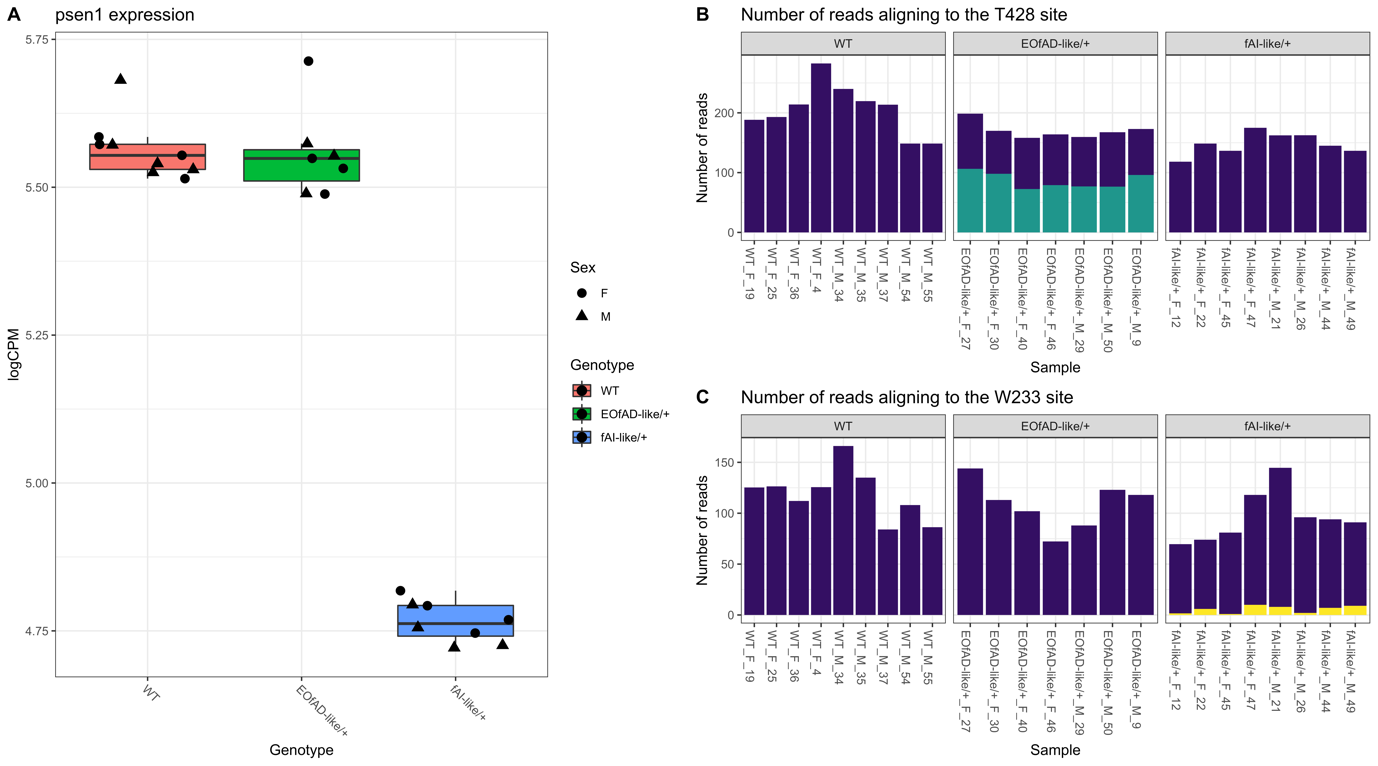

**Figure S7:** **A** Overall expression of *psen1* in 6 month old zebrafish brains in log2 counts per million (logCPM). **B** The number of reads aligning to the GRCz11 genome at the T428 site and **C** the W233 site in *psen1.* Bars appear purple if aligning to the wild type allele, blue if the aligned reads contained the T428del mutation, and yellow if the aligned reads contained the W233fs mutation. For each sample, the number of reads was calculated as the average number of reads over the three nucleotides deleted for the T428 site, and the two nucleotides affected at the W233 site.

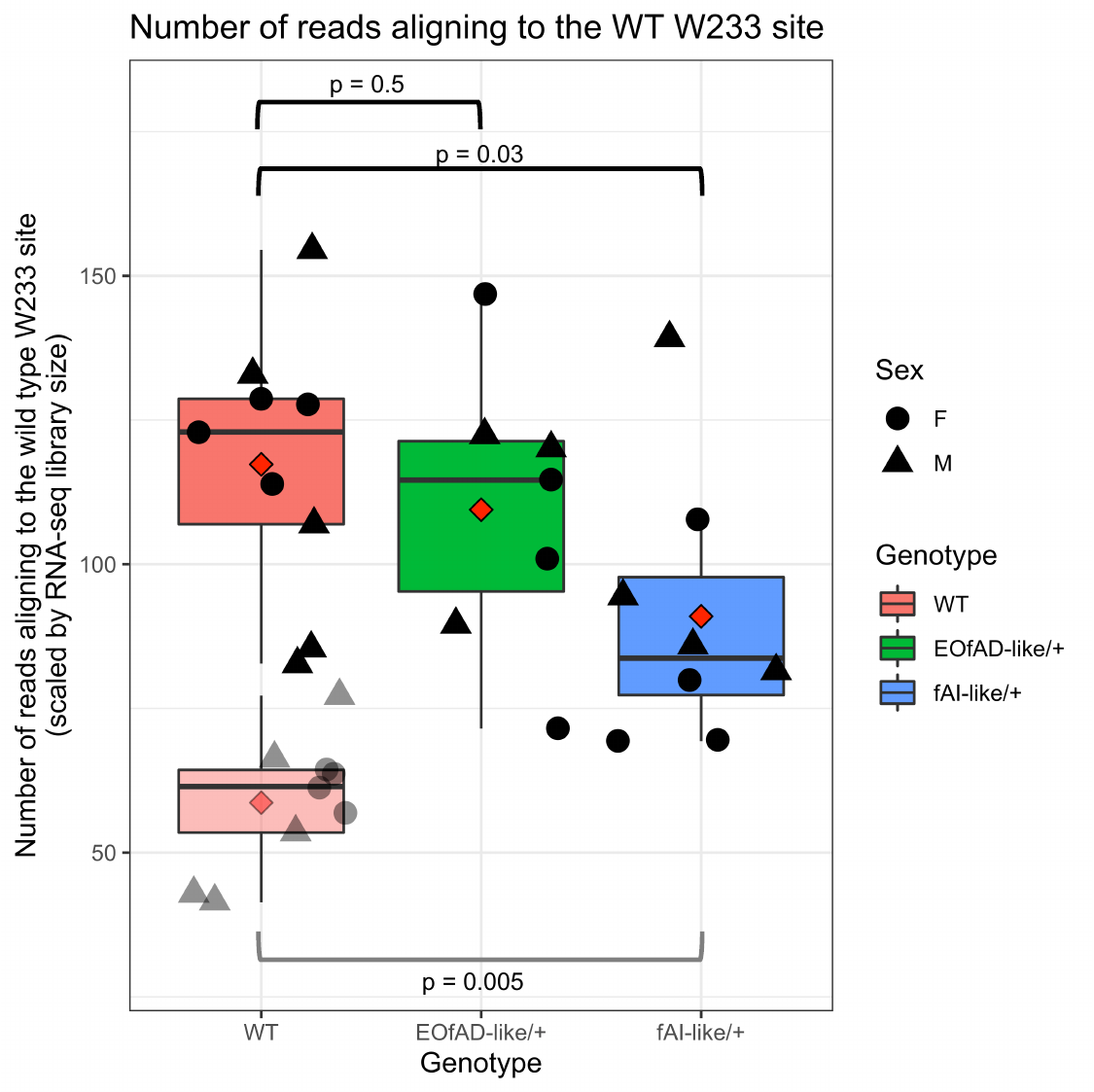

**Figure S8:** For each sample, the number of reads was calculated as the average number of reads over two nucleotides affected at the W233 site in *psen1*, then scaled for RNA-seq library size (TMM method). Pairwise comparisons between genotypes for these measures (indicated by the red diamonds) was performed using Welch’s two sample t-tests. To investigate whether the expression of the wild type *psen1* allele in fAI-like/+ brains was greater than 50% of the expression of the wild type allele in wild type brains, the scaled reads for wild type brains were divided by two (shown in the graph by the more transparent points), then these were subject to Welch’s two sample t-test.

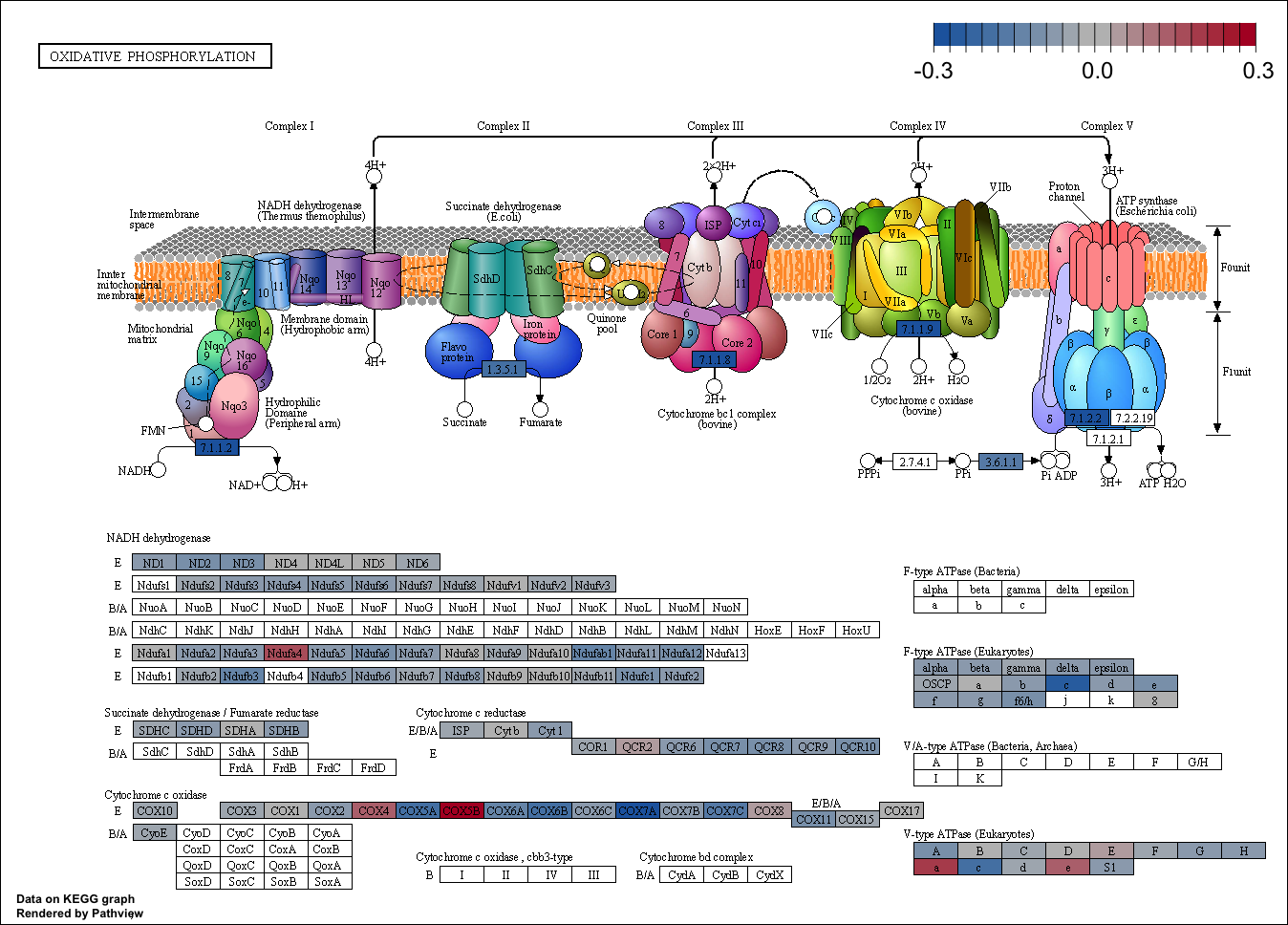

**Figure S9:** Pathview visualisation of the *KEGG_OXIDATIVE_PHOSPHORYLATION* gene set. Colour of the cells indicates the logFC in EOfAD-like/+ brains.

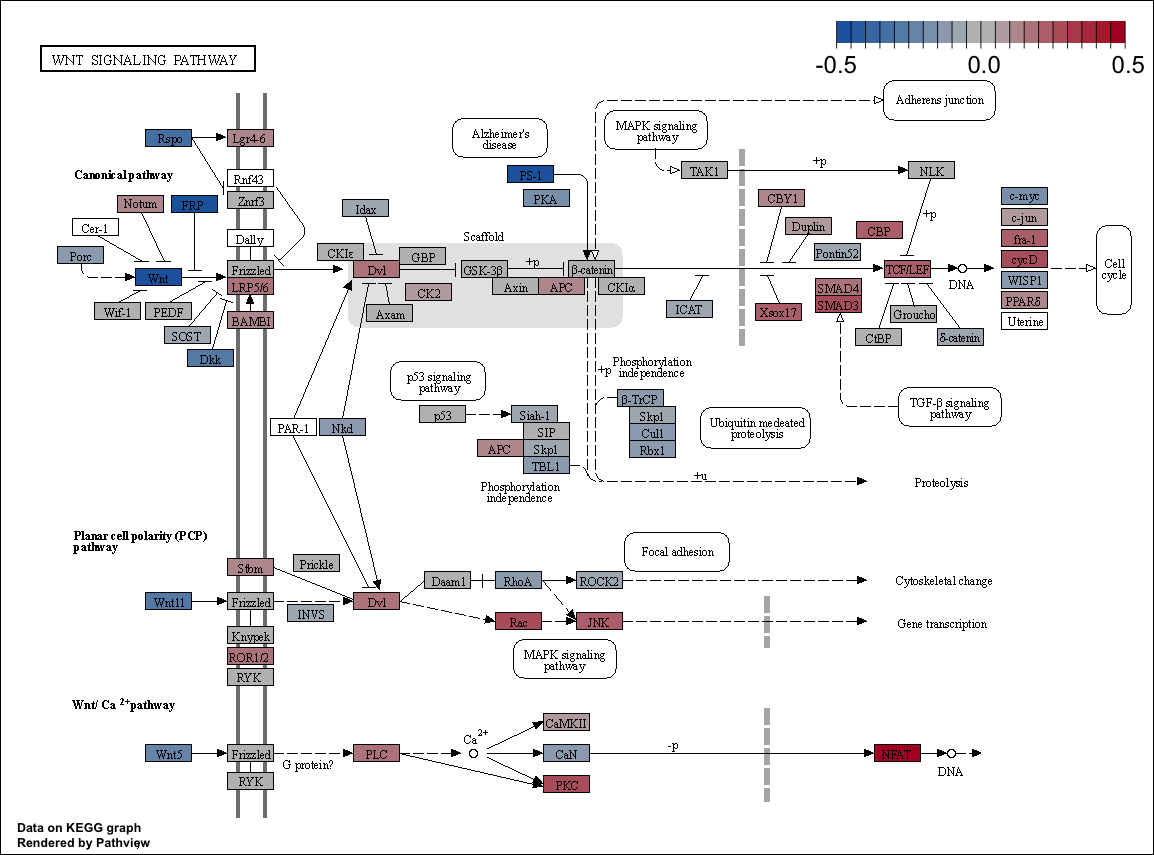

**Figure S10:** Pathview visualisation of the *KEGG_WNT_SIGNALING_PATHWAY* gene set. Colour of the cells indicates the logFC in fAI-like/+ brains.

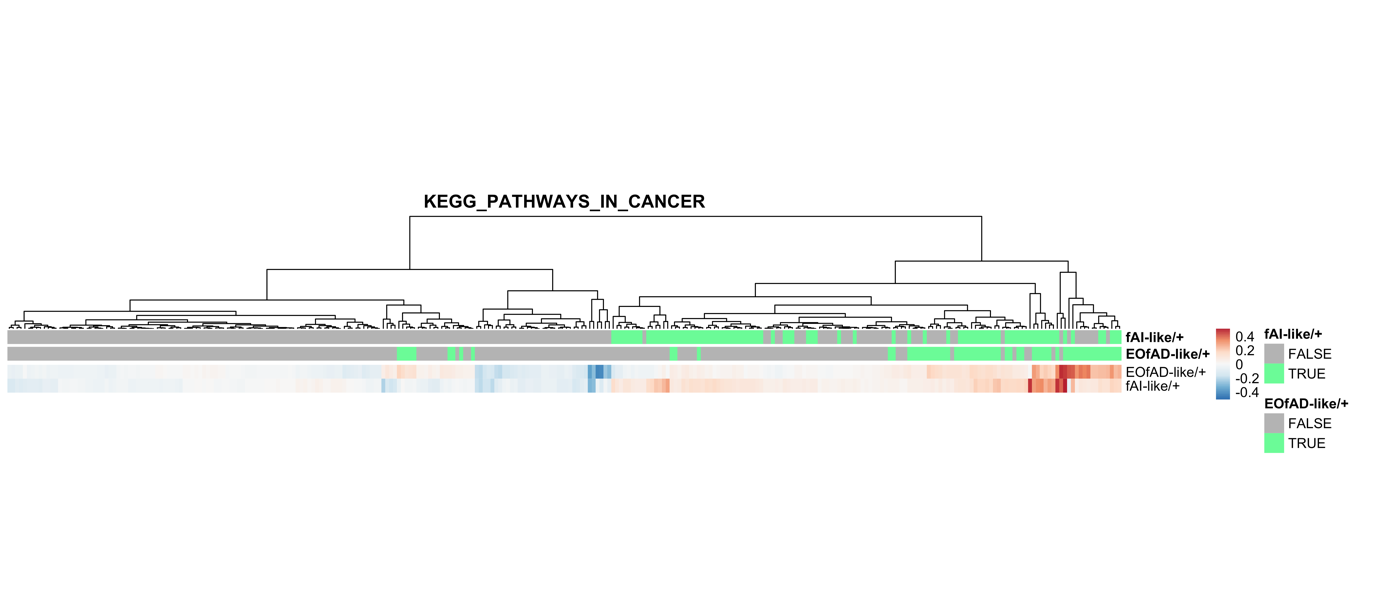

**Figure S11:** Heatmap indicating the logFC of genes in the *KEGG_PATHWAYS_IN_CANCER* gene set in EOfAD-like/+ and fAI-like/+ brains. Cells in the heatmap are labelled in green if found in the leading edge from the fgsea algorithm and grey if they were not.

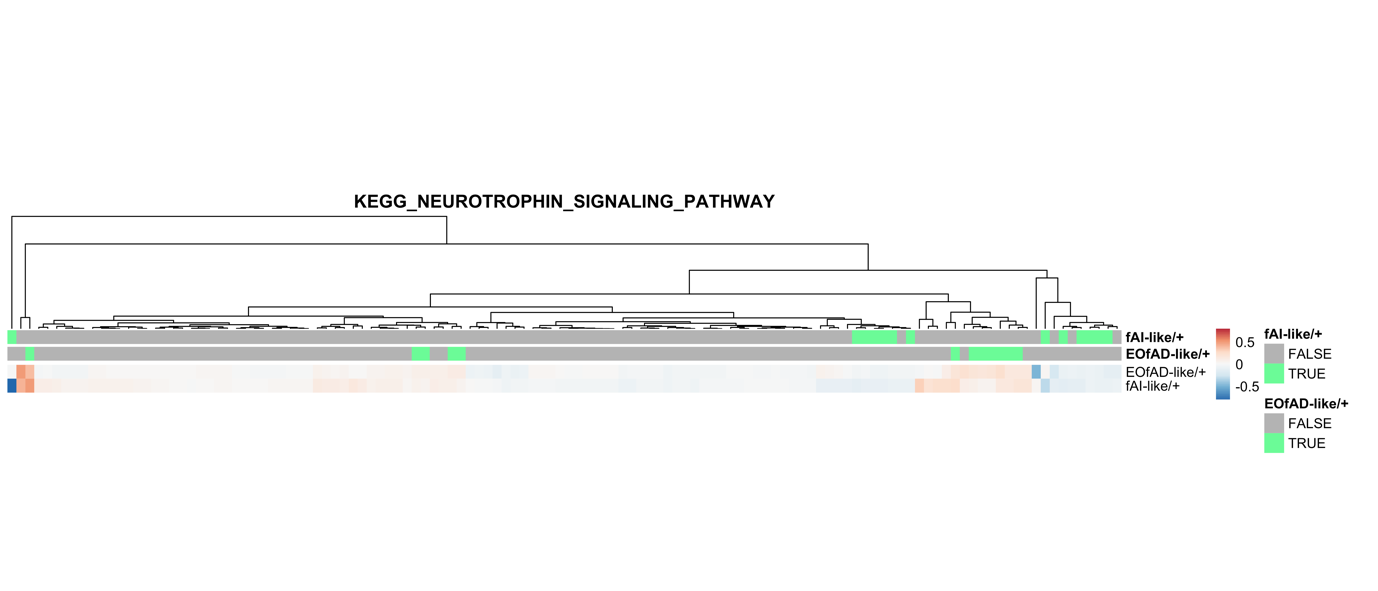

**Figure S12:** Heatmap indicating the logFC of genes in the *KEGG_NEUROTROPHIN_SIGNALING_PATHWAY* gene set in EOfAD-like/+ and fAI-like/+ brains. Cells in the heatmap are labelled in green if found in the leading edge from the fgsea algorithm and grey if they were not.

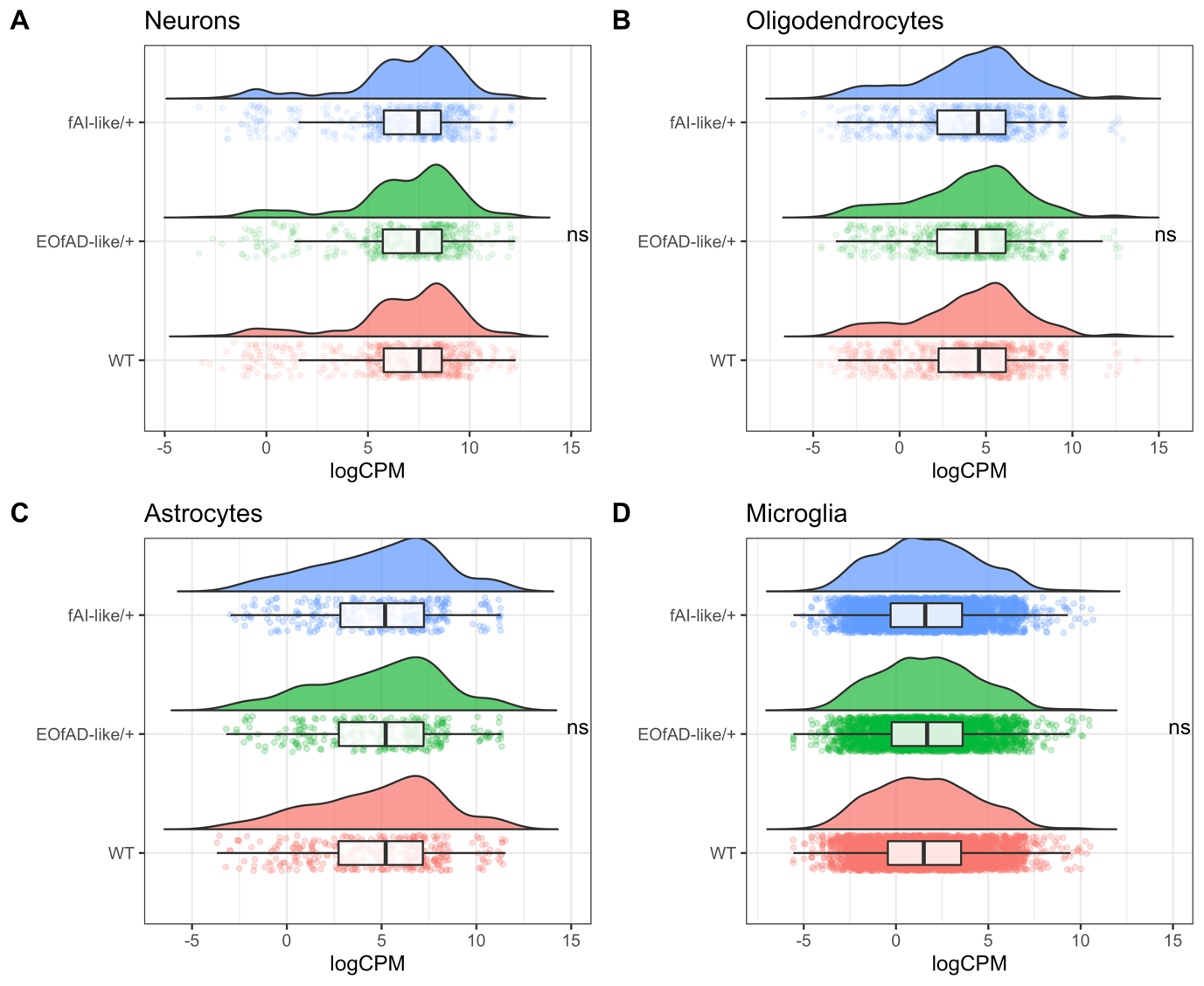

**Figure S13:** Cell type proportions appear to be consistent across samples. The raincloud plots below summarise the expression (logCPM) of marker genes for four broad cell types in zebrafish brains: neurons, oligodendrocytes, astrocytes and microglia. The marker genes for **A** neurons, **B** oligodendrocytes and **C** astrocytes were obtained from [1], while the **D** microglia markers were obtained from [2]. No significant differences (ns) of expression of these markers was found across genotypes using *fry* with a directional hypothesis, supporting that cell type proportions are consistent.
